# Surface protein profiling of milk and serum extracellular vesicles unveil body fluid and cell-type signatures and insights on vesicle biogenesis

**DOI:** 10.1101/2022.10.24.513472

**Authors:** Alberta Giovanazzi, Martijn J.C. van Herwijnen, Gerbrich N. van der Meulen, Marca H.M. Wauben

## Abstract

The promise of extracellular vesicles (EVs)-based liquid biopsy resides in the identification of specific signatures of EVs of interest. Knowing the EV profile of a body fluid can facilitate the identification of EV-based biomarkers of diseases. To this end, we characterised purified EVs from paired human milk and serum by surface protein profiling of cellular markers in association with gold standard EV markers (tetraspanins CD9, CD63 and CD81). By using the MACSPlex bead-based flow-cytometry assay with pan-tetraspanin detection (i.e. simultaneous CD9, CD63 and CD81 detection), besides specific breast epithelial cell signatures in milk EVs and platelet signatures in serum EVs, we also identified body fluid-specific markers of immune cells and stem cells. Interestingly, comparison of pan-tetraspanin and single tetraspanin detection unveiled both body fluid-specific tetraspanin distributions and specific tetraspanin distributions associated with certain cellular markers, which were used to model the potential biogenesis route of different EV subsets and their cellular origin.

## Introduction

Extracellular vesicles (EVs) are submicron lipid bilayer-delimited particles naturally released by cells that act as mediators of inter-cellular communication by targeting biologically active molecules to adjacent and distant cells^1^. Cells in body tissues communicate by releasing EVs into proximal body fluids, such as breast milk and blood^2^. Circulating EVs can originate from cells present in the body fluids, cells lining the cavities of extruded fluids or from tissue-resident cells^2^, and for this reason they can carry body fluid-specific and cell-specific signatures. Additionally, the molecular make-up of EVs can be affected by the status of their originating cells and, as such, EVs can be enriched or depleted for specific surface proteins, resulting in specific protein biomarker profiles associated with (patho)physiological conditions^3,4^. Nowadays, EV-based biomarker discovery attracts a lot of attention for monitoring disease and health status. The promise of EV-based biomarkers resides in the specific combination of different EV molecules resulting in a “combined” biomarker that outperforms single component-based biomarkers. The discovery of such EV-specific “combined” profiles is strongly dependent on the purification of EVs from the body fluid. Since body fluid components differ in colloidal properties^5^ and non-EV particles with overlapping characteristics of EVs^6^ are present in body fluids, this complicates EV isolation and downstream analysis. For example, lipoproteins in blood and casein micelles in milk, which co-isolate to various degrees with EVs^7,8^, can act as confounders in (semi)-quantitative particle analyses such as multiplex assays.

To identify EV-based biomarkers of disease or disturbed homeostasis, knowing the “normal” molecular profile of EVs in different body fluids is also of utmost importance. The tetraspanins CD9, CD63 and CD81 are considered *bona fide* EV-associated markers and are commonly used for “total” EV detection in immunoassays. CD9, CD63, and CD81 are the three most-studied tetraspanins in the EV field due to their primary functions in EV formation, cargo selection/sorting and EV release and uptake^9^. Via their extracellular domains, tetraspanins associate with other tetraspanins and surface proteins, forming “tetraspanin webs”^10^. Importantly, the association of tetraspanins with a multitude of proteins results in a variety of surface protein profiles. Moreover, in recent years, it has been reported that the distribution of tetraspanins is more heterogeneous than assumed across single EVs^11–14^. Importantly, specific combinations of CD9, CD63 and/or CD81 on EVs can give information on their specific biogenesis route^11,12^ and their associations with specific EV surface proteins can give clues on their cell of origin. Consequently, the commonly used pan-tetraspanin immunodetection in multiplex protein profiling can mask important biological information and can even hamper the identification of EV-based “combined” protein biomarkers.

In the present study, we studied surface protein profiles of milk and serum EVs, purified with validated methods for minimal body fluid contamination^15–18^, by performing multiplexed flow cytometric analysis (MACSPlex) based on pan-tetraspanin EV detection and individual tetraspanin detection. Using this approach we were able to identify body fluid and cell-type specific EV-associated protein profiles and gained insights on possible biogenesis routes of EV subsets.

## Results

### Concentration and size determination of purified EVs from paired human milk and serum samples

For comparative analysis, EVs were isolated from paired human milk and serum samples, both donated on the same day by non-allergic and allergic mothers. Since equal amounts of EVs are crucial for reliable comparisons^19^, robust quantitative EV analysis was required. By using our optimized EV isolation protocol, consisting of differential centrifugation without high-*g* pelleting (>10.000xg), known to cause blood-derived EV aggregation^20,21^, followed by density gradient separation and size exclusion chromatography, non-EV colloidal structures present in blood or milk that can outnumber EVs by orders of magnitudes and, as such, hamper quantitative EV analysis, were largely depleted from the samples^15,18^. Subsequently, EV concentration and size were determined by Nanoparticle Tracking Analysis (NTA). For reliable analyses, only measurements with 10-110 particles in frame and at least 1000 valid tracks per EV sample **(Supplementary Fig. S1)** were processed^22^. No differences in particle concentrations and size distributions were observed between EVs derived from non-allergic and allergic donors, however, these parameters differed between EVs present in milk and in serum. The concentration of milk EVs isolated from n=9 individual donors ranged from 5×10^10^ - 1.2×10^11^ particles/ml (average ± standard deviation = 8×10^10^ ± 2.8×10^10^) while the concentration of serum EVs ranged from 5×10^9^ - 1.5×10^10^ particles/ml (average ± standard deviation = 1×10^10^ ± 4.3×10^10^) **(Fig. 1a)**. The modal size of milk EVs and serum EVs was in the range of 230 ± 13 nm and 105 ± 10 nm, respectively **(Fig. 1b)**. Size distribution histogram overlays of paired milk and serum EV samples show this size difference for every individual donor **(Fig. 1c)**. Overall, the milk EV samples had significantly higher particle concentration and bigger size than the paired serum EV samples **(Fig. 1d)**.

**Figure 1.**
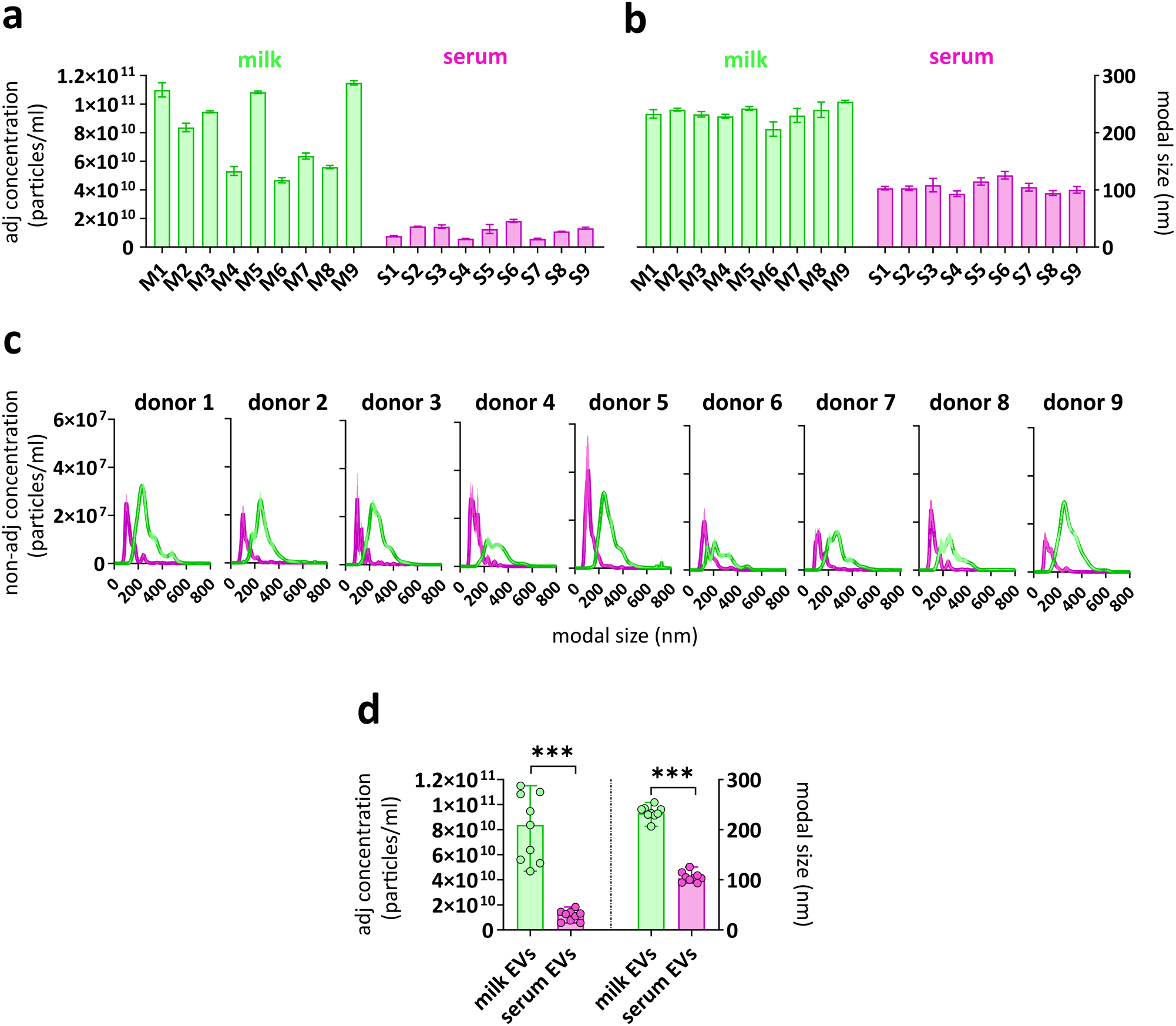
Concentration and size distribution of paired milk EV and serum EV samples. NTA was used to determine **(a)** the adjusted nanoparticle concentration (particles/ml) and **(b)** the modal size (diameter) of n=9 milk EV and paired serum EV samples. Bars represent mean ± SD from three captures of 60 seconds per sample. **(c)** NTA histograms show the non-adjusted concentration and the size distribution of paired milk EVs (green) and serum EVs (pink). **(d)** The median adjusted nanoparticle concentration and modal size of n=9 paired milk EV samples and serum EV samples. Significance was tested by 2-tailed non-parametric Mann-Whitney T test, p<0.001 (***).

### Optimization of the MACSPlex assay analysis for comparative surface protein profiling of purified milk and serum EVs

To study the milk and serum EV surface proteins, the multiplex bead-based flow cytometry MACSPlex platform was used. The principle of the assay relies on the use of hard-dyed bead populations, each coupled to different antibodies that recognize 37 potential EV surface antigens (and 2 internal isotype controls)^19,23^. Beads are distinguished from each others by their respective fluorescence characteristics (FITC VS PE) and bound EVs are detected by a cocktail of APC-conjugated antibodies against the gold standard EV markers CD9, CD63 and CD81 (Pan-tetraspanin detection) (APC VS FITC) **(Supplementary Fig. S2)**.

To perform a robust and comparative analysis, we first defined the optimal input dose of EVs by testing 5 different amounts, ranging from 10^8^ to 10^9^ EVs purified from milk and 5*10^7^ to 5*10^8^ EVs purified from serum **(Supplementary Table S2)**. As expected, the percentage of APC^+^ events proportionally increased with increasing input numbers of EVs **(Fig. 2a, Supplementary Fig. S2)**. When comparing the percentage of total APC^+^ events between the same amounts of serum and milk EVs, no significant differences were observed, confirming that same input dose of EVs from the two body fluids was used **(Supplementary Fig. S2)**. Additionally, the signal intensities for the tetraspanin CD9, CD63 and CD81 capture bead populations, but not for the internal isotype control bead populations, increased with increasing EV amounts **(Fig. 2b and Supplementary Fig. S2)**. The median APC signal of the three tetraspanins increased by approximately 10X from the lowest to the highest EV dose tested and thus showed a proportional titration for both serum and milk samples **(Fig. 2c)**. Since 5*10^8^ was the highest dose of serum EVs that could be obtained without further sample concentration and which resulted in robust and clear APC detection signals, we selected 5*10^8^ input EVs for further comparative analyses of milk and serum EVs. Overall, these results showed that the MACSPlex platform is suitable for comparative analysis of EVs purified from human milk and serum.

**Figure 2.**
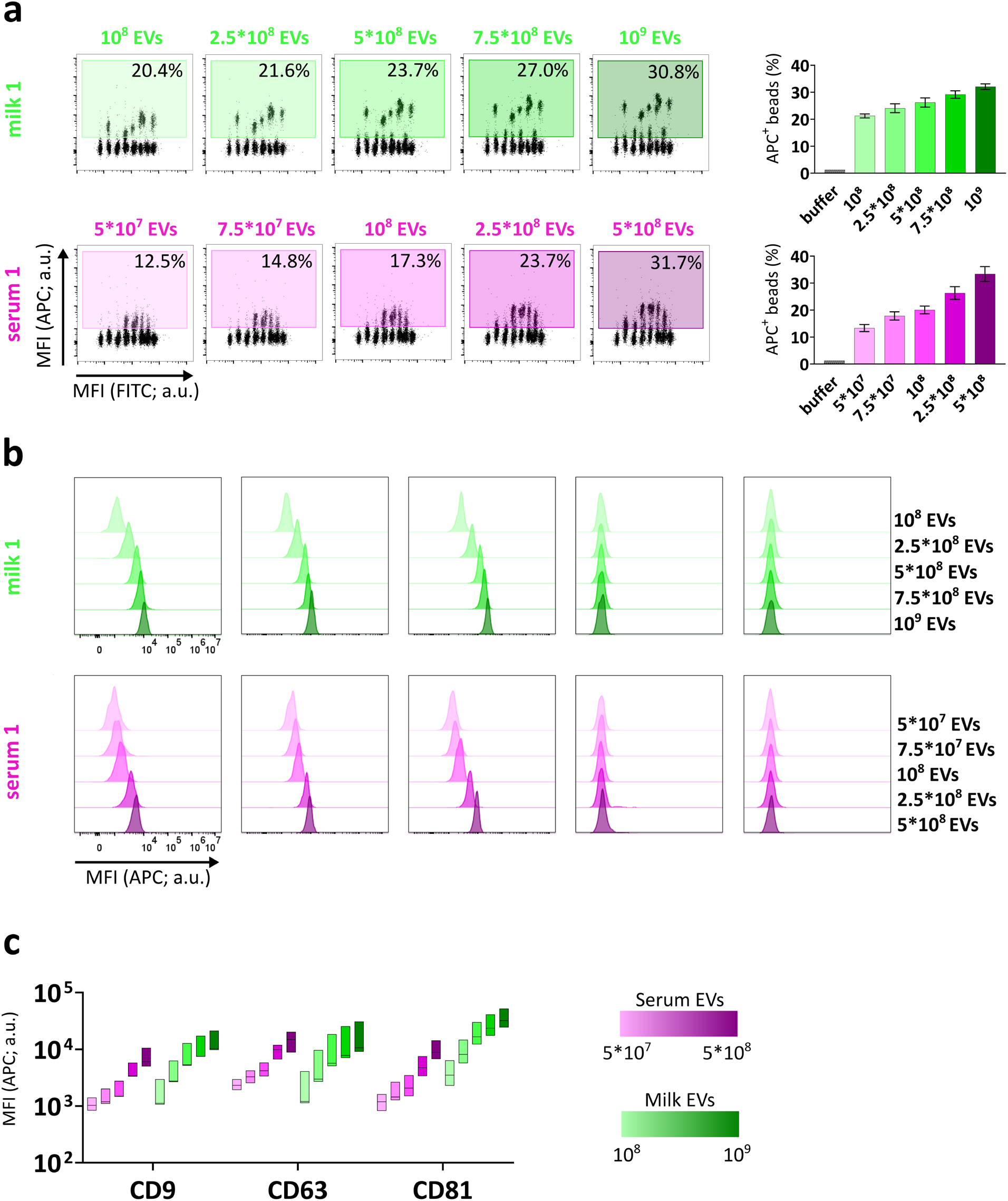
Titration of APC detection signals is proportional to input EV numbers. **(a)** Flow cytometry dot plots of five input amounts of milk EVs (green) and serum EVs (pink) from donor 1 depicting the percentage of APC^+^ beads (relative to single beads). The bar graphs show the percentage of APC^+^ beads as mean ± SD of three donors (donor 1, donor 2, donor 3). **(b)** Flow cytometry histograms representing MFI APC signal relative to CD9, CD63, CD81 capture beads and isotype control REA and mIgG1 in relation to increasing doses of milk EVs (green) and serum EVs (pink) from donor 1. **(c)** The floating bar graph depicts the median + min and max values (n=3 donors) of MFI APC relative to CD9, CD63, CD81 capture beads at different EV doses. a.u. = arbitrary unit.

### Pan-tetraspanin EV detection unveil body fluid-specific protein signatures

Next, we characterized the surface protein profiles of milk and serum EVs detected with pan-tetraspanin antibody cocktail **(Supplementary Table S3)**. Proteins whose Median Fluorescent Intensity of the APC signal (MFI APC) was lower than the respective isotype controls were considered non-detected **(Fig. 3a and Supplementary Table S4)**. Based on this criterium, 15 and 17 proteins were detected on EVs from milk and serum, respectively **(Fig. 3b)**, with a total number of 21 detected proteins, and no differences were observed between EVs from non-allergic donors (n=5) and allergic donors (n=4) **(Supplementary Fig. 4)**.

**Figure 3.**
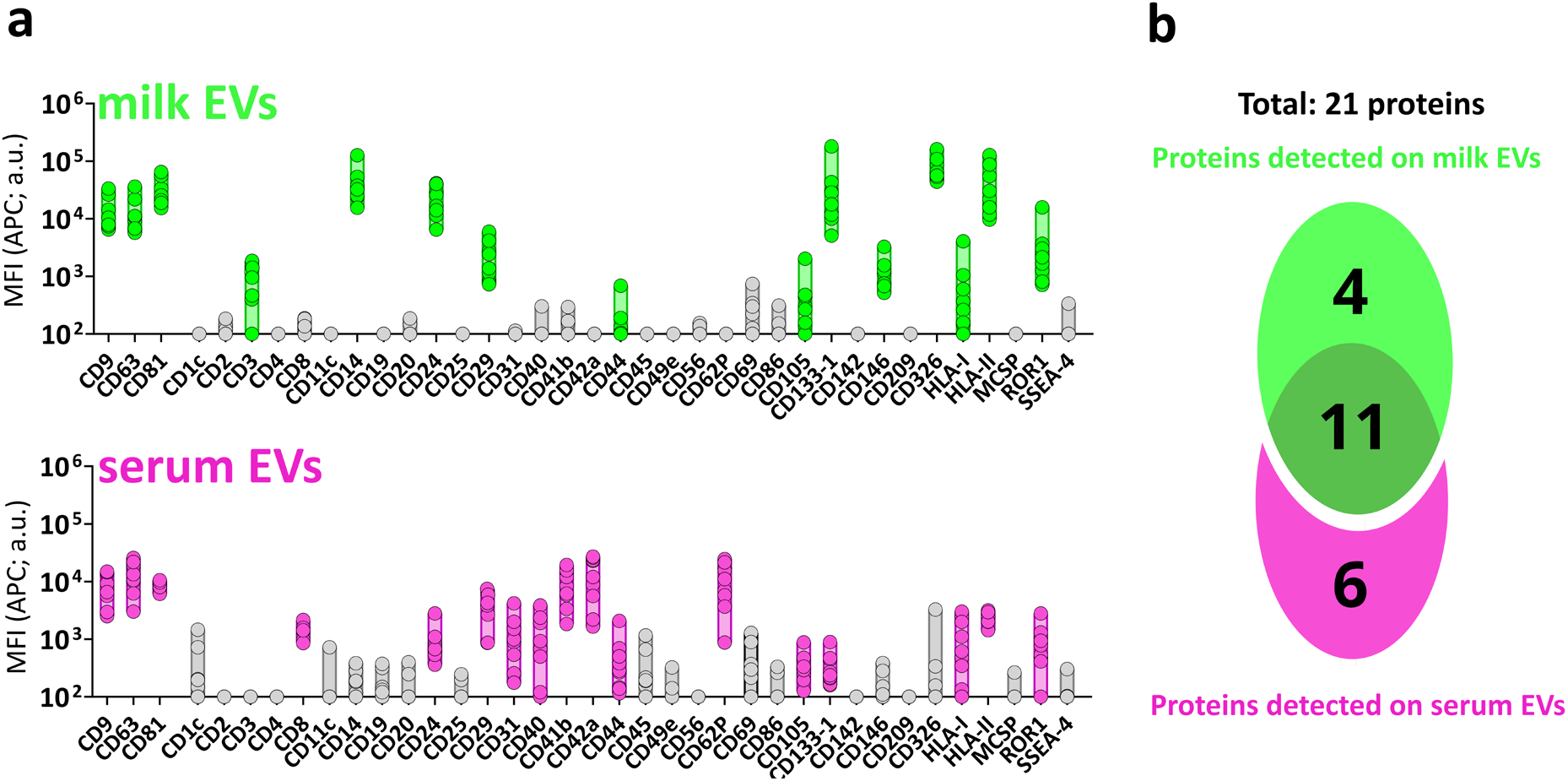
Protein distribution on milk EVs and serum EVs with pan-tetraspanin detection. **(a)** MFI APC signal of pan-tetraspanin detection relative to the 37 MACSPlex proteins on n=9 paired milk EV (green) and serum EV (pink) samples (5*10^8^ EVs). In grey proteins whose MFI was not significantly higher than the MFI of the corresponding internal isotype control (non detected). Significance was determined by non-parametric Kruskal-Wallis Test with post-hoc Dunn’s multiple comparison (Supplementary Table S4), p≥0.05 (ns), p<0.05 (*), p<0.01 (**), p<0.001 (***). **(b)** Venn Diagram depicting unique and common proteins of milk EVs and serum EVs. a.u. = arbitrary unit.

11 out of these 21 detected proteins were in common between milk and serum EVs and no significant differences were observed in the MFI APC pan-tetraspanin signals of 7 proteins (CD9, CD63, HLA-I, CD29, CD44, CD105, ROR1) **(Supplementary Fig. S3)**. The flow cytometry dotplots representing the MFI APC of the pan-tetraspanin detection of the remaining 14 detected proteins show different profiles in milk EVs and serum EVs (**Fig. 4a** **and** **4b**). Specifically, the pan-tetraspanin detection levels of CD326, CD14, HLA-II, CD133-1, CD81, CD24, CD146 and CD3 were significantly higher in milk EVs, and 4 proteins were exclusively detected on milk EVs (CD326, CD14, CD146 and CD3) **(Fig. 4c)**. The pan-tetraspanin signals for CD42a, CD62P, CD41b, CD31, CD40, CD8 were only detected in serum EVs and undetectable in milk EVs **(Fig. 4d)**. Taken together, these results show that besides common surface proteins, milk EVs and serum EVs also express body fluid-specific surface proteins.

**Figure 4.**
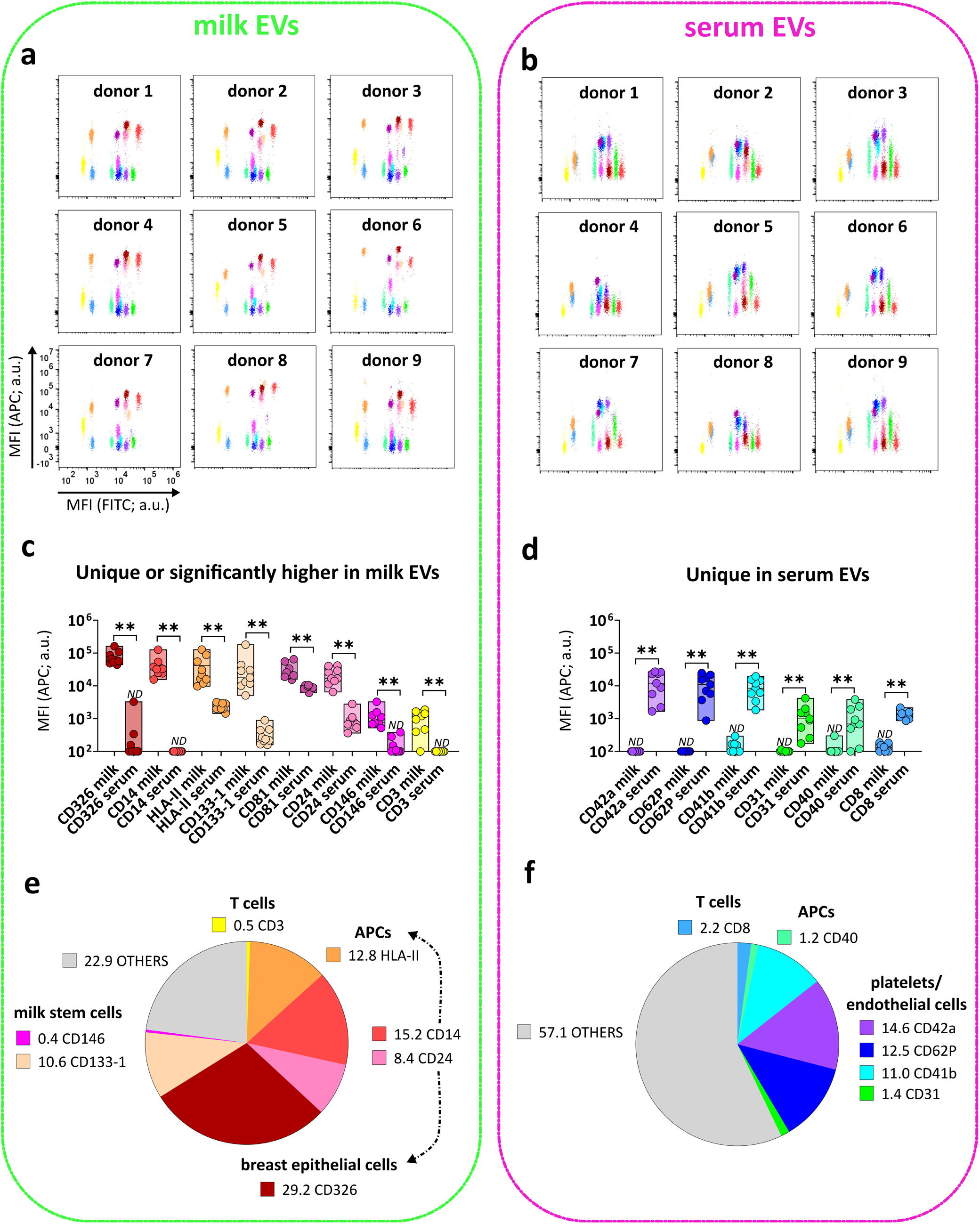
Surface proteins unique or significantly differently expressed on milk EVs and serum EVs. Flow cytometry dotplots from milk EVs **(a)** and serum EVs **(b)** representing all the 14 proteins enriched either in milk EVs (8 proteins) **(c)** or in serum EVs (6 proteins) **(d)**. Bars indicate minimum and maximum value with median of MFI APC pan-tetraspanin values from n=9 EV samples per body fluid. Statistical significance was determined by multiple non-parametric matched pairs Wilcoxon test with Benjamini and Hochberg FDR method, p<0.01 (**). Percentage expression of the total MFI APC pan-tetraspanin signal (MFI APC values of each capture bead population was converted into a percentage of the total signal) for proteins enriched in milk EVs **(e)** and in serum EVs **(f).** Proteins are clustered based on known cellular markers. “Others” (indicated in grey) includes protein signals lower than the other body fluid (e.g. CD133, HLA-II, CD81, CD24 for serum EVs), non-significantly different proteins (Supplementary Fig. S3a) and proteins below the isotype threshold (Fig. 3). ND is non-detectable as defined by the isotype threshold (Fig. 3). a.u. = arbitrary unit.

Next, we attempted to identify the potential cells of origin of the body fluid-specific/enriched EV proteins present in milk and serum (**Fig. 4e** **and** **4f)**. In milk EVs, 11% of the total MFI APC signal of the pan-tetraspanin detection in the MACSPlex array was derived from CD133-1 (10.6%) and CD146 (0.4%), which are associated with breast milk stem cells^24–27^, 29.2% from CD326, a marker for luminal breast epithelial cells (lactocytes or milk secretory cells)^8,28^, and 12.8% from the antigen-presenting cells (APCs) marker MHC-II. CD14 (15.2%) and CD24 (8.4%) were also present on milk EVs, which could reinforce the hypothesis that milk carries a population of EVs derived from antigen-presenting cells. However, similarities exist between certain leukocytic and breast epithelial cell populations^29,30^. For example, CD14, despite being first identified as marker of mature monocytes, can be associated to mammary epithelial cells^31^. Similarly, CD24 has often been associated to B cells. However, in the context of breast milk, CD24 most likely marks luminal epithelial cells^32^. Additionally, the glycoprotein CD44^25,33^ and Integrinβ1/CD29^25^ are reported to be widely expressed on basal myoepithelial breast cells (from the ducts and alveoli of mammary gland). CD44 has also been described as a marker of stem cells in milk^26^. Importantly, CD44 is a rather general marker for many different cells making it difficult to define the cell of origin of EVs expressing CD44. Albeit much lower, CD3 was uniquely detected on milk EVs, as 0.5% of total MFI APC pan-tetraspanin signal, suggesting the origin of CD3-expressing milk EVs from T cells. Overall, these data showed that the majority of milk EVs (77.1% of the total MFI APC pan-tetraspanin signal of the array) carry features hinting at their possible origin from mammary epithelium cells, as well as stem cells, APCs and T cells **(Fig. 4e)**.

In serum EVs, the most prominent signal accounted for platelets-associated markers, i.e. platelet-derived glycoprotein CD41b and platelet glycoprotein IX CD42a, respectively 11% and 14.6% of the total MFI APC pan-tetraspanin signal in the array. 13.9% of the signal reflected the ongoing platelet activation and interaction with endothelium (P-selectin CD62P and platelet endothelium adhesion molecule PECAM-1/CD31)^34^, which suggests that EVs derived from activated platelets, as expected, are highly abundant in serum **(Fig. 4f)**. Besides platelet signatures in serum EVs, serum-specific proteins corresponding to antigen presenting cells (i.e. CD40 1.2% of total MFI APC pan-tetraspanin signal) and T cells (i.e. CD8 2.2% of total MFI APC pan-tetraspanin signal) were detected. Overall, the serum-specific proteins make up 42.9% of the total MFI APC pan-tetraspanin signal of the array.

### Tetraspanins are heterogeneously distributed on EVs from milk and serum

In the pan-tetraspanin detection, the total signals of CD9, CD63 and CD81 tetraspanins were used to detect EVs. However, tetraspanins are unequally distributed on EV subsets^11,12^, as also demonstrated in the pan-tetraspanin detection for the CD81 capture bead showing significantly higher signals in milk EVs compared to serum EVs, both in the group analysis **(Fig. 4c and Supplementary Fig. 5)**, as well as in the paired analysis of individual donors **(Supplementary Fig. 5)**. These data demonstrated that the distribution of tetraspanins on EVs is not homogeneous and can differ between EVs from various body fluids. To explore EV subset differences between milk and serum in greater detail, we next compared pan-tetraspanin detection (αCD9, αCD63 and αCD81 antibody cocktail) with single tetraspanin detection (αCD9, αCD63 or αCD81 single antibody). While the CD9 capture bead signal based on pan-tetraspanin detection was not significantly different between serum EVs and milk EVs **(Supplementary Fig. 5)**, the single tetraspanin detection showed that the main contributor to the CD9 bead signal in the pan-tetraspanin detection differed between milk and serum; i.e. CD81 in milk (pan-t VS αCD81, p≥0.05, ns) and CD9 in serum (pan-t VS αCD9, p≥0.05, ns) **(Fig. 5a and Supplementary Fig. 6)**. A similar pattern was observed for the CD63 capture bead signal: while no significant differences in the pan-tetraspanin detection were observed between serum and milk EVs **(Supplementary Fig. 5)**, CD81 in milk (pan-t VS αCD81, p≥0.05, ns) and CD9 in serum (pan-t VS αCD9, p≥0.05, ns) were the main contributors to the pan-tetraspanin CD63 capture bead signal **(Fig. 5b and Supplementary Fig. 6)**. In **Figure 4c and Supplementary Fig. 5**, we showed a significant difference between the pan-tetraspanin detection of CD81 capture bead of milk EVs and serum EVs, with a significantly higher signal in milk EVs. Also for the CD81 capture bead signal, the main contributor was CD81 in milk EVs (pan-t VS αCD81, p≥0.05, ns) and CD9 in serum EVs (pan-t VS αCD9, p≥0.05, ns) **(Fig. 5c and Supplementary Fig. 6)**. In conclusion, the comparison of pan-tetraspanin and single tetraspanin detection revealed body fluid-specific contributions of individual tetraspanins to the pan-tetraspanin signal of the three tetraspanin capture bead populations (i.e. in serum CD9 dominated the pan-tetraspanin signal of CD9, CD63 and CD81 capture bead populations, while, in milk, CD81 was the most prominent).

**Figure 5.**
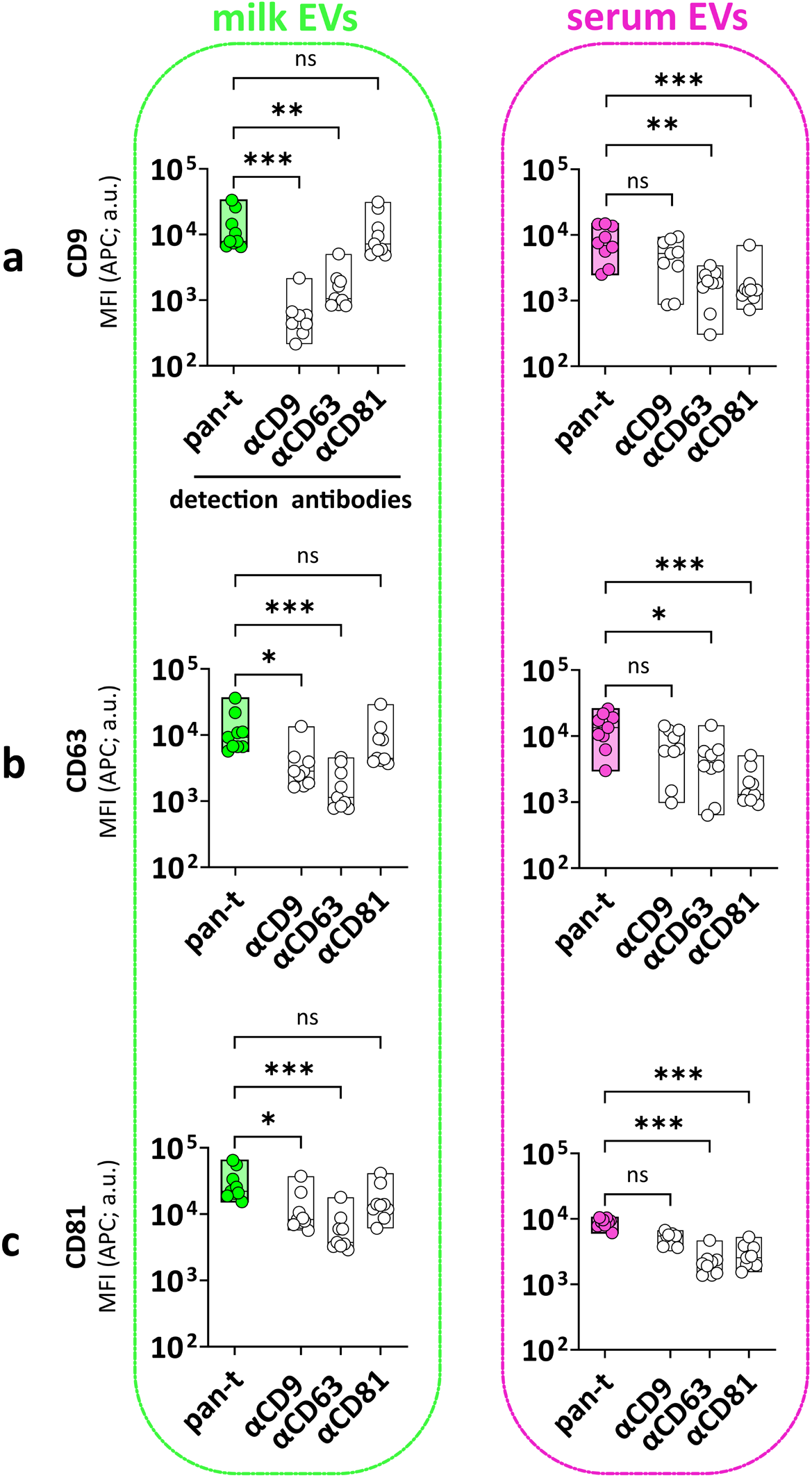
Heterogeneous distribution of tetraspanins on EVs from milk and serum. MFI APC of CD9 **(a)**, CD63 **(b)** and CD81 **(c)** capture bead populations (x-axis) detected with standard pan-tetraspanin antibody (pan-t: αCD9, αCD63 and αCD81) and single tetraspanin antibody (αCD9, αCD63 or αCD81) (y-axis). Bar graphs delimited by minimum and maximum value with median of MFI APC values from n=9 EV samples per body fluid. Non-parametric Kruskal-Wallis Test with post-hoc Dunn’s multiple comparison was performed to compare pan-tetraspanin with each single tetraspanin detection, p≥0.05 (ns), p<0.05 (*), p<0.01 (**), p<0.001 (***). a.u. = arbitrary unit.

### Single tetraspanin detection of milk EVs and serum EVs unveiled different tetraspanin profiles depending on the capture antibody

To obtain more insight into different EV subpopulations in milk and serum, we analysed the single tetraspanin detection signals of milk and serum EVs bound to the capture bead populations identified in **Figure 3** (**Supplementary Figure 7** shows the MFI APC signals from the three single tetraspanin detection antibodies). Next, we compared the MFI APC pan-tetraspanin detection signals with single tetraspanin signals (**Fig. 6a** **and** **6b**). For milk EVs **(Fig. 6a)**, we identified 7 proteins/capture bead populations (i.e. CD326, CD14, HLA-II, CD24, CD29, ROR-1 and CD44) for which the MFI APC pan-tetraspanin detection signal was comparable to single CD81 detection signal (pan-t VS αCD81, p≥0.05, ns), suggesting that the signal from CD81 detection was the main contributor to the pan-tetraspanin signal of EVs bound to these capture bead populations. For 6 out of these 7 bead populations (i.e. CD326, CD14, HLA-II, CD24, CD29 and ROR-1), single CD9 and single CD63 detection signals were significantly lower than the pan-tetraspanin detection. In contrast, for the CD44 capture bead the detection signals from milk EVs by single CD9 and single CD63 were below the isotype threshold, indicating that CD44-positive milk EVs only expressed CD81. For 3 other milk EV markers (i.e. CD133-1, CD146 and HLA-I) single CD9 and single CD81 detection contributed equally to the pan-tetraspanin signal. In contrast to CD133-1 and CD146, milk EVs captured by HLA-I did not express CD63 (below the isotype threshold). Finally, the MFI APC pan-tetraspanin signal of milk EVs captured by CD3 and CD105 was comparable to single CD63 and CD81 detection, while these EVs did not express CD9 (below the isotype threshold).

**Figure 6.**
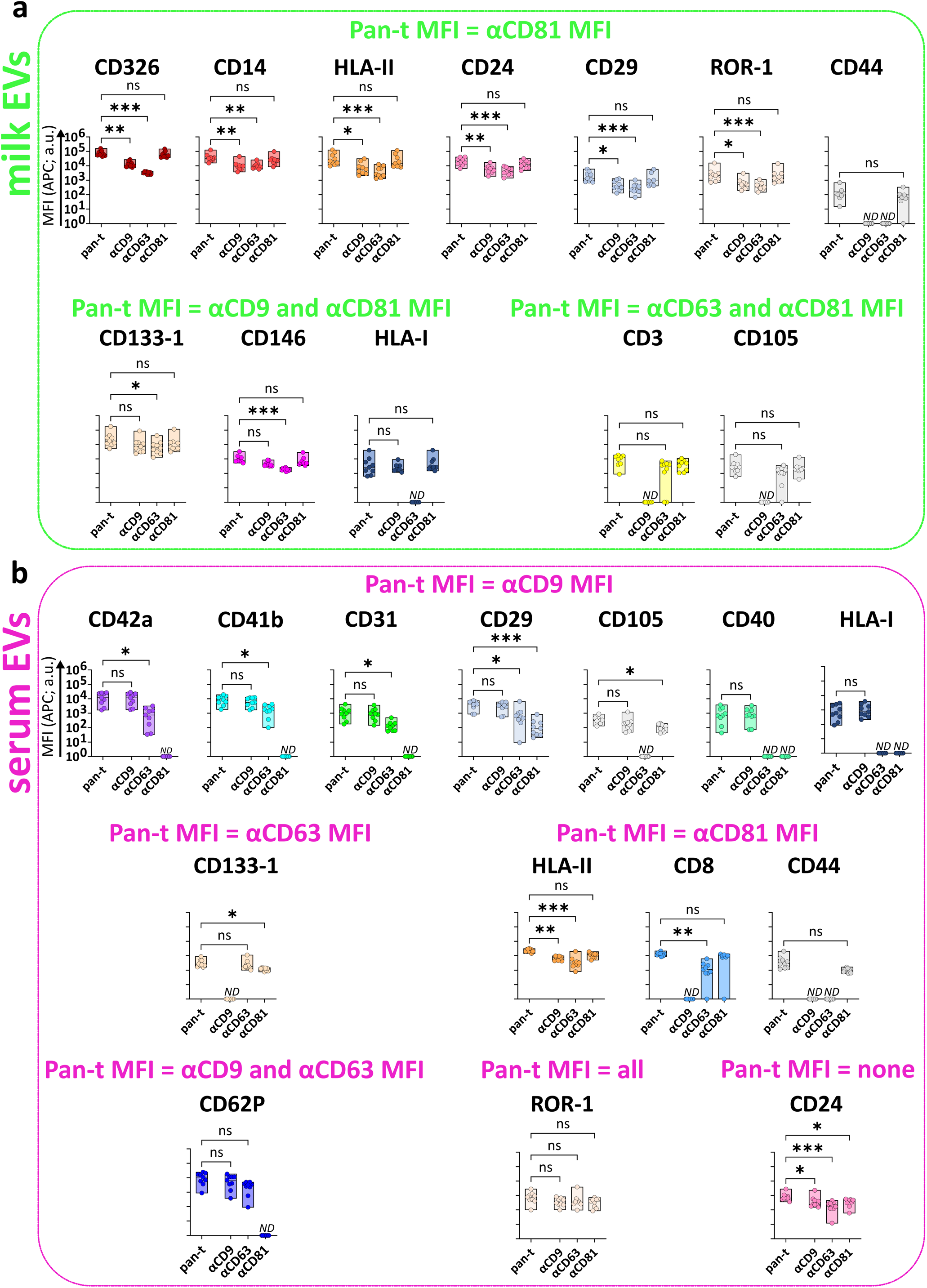
Comparison of pan-tetraspanin and single tetraspanin detection of milk EVs and serum EVs bound to different protein capture bead populations. MFI APC signals of capture bead populations (identified in Figure 3) on milk EVs **(a)** and serum EVs **(b)** from pan-tetraspanin detection (pan-t: αCD9, αCD63 and αCD81) in comparison to each individual tetraspanin detection (αCD9, αCD63 or αCD81). Bar graphs delimited by minimum and maximum value with median of MFI APC values from n=9 EV samples per body fluid. Non-parametric Kruskal-Wallis Test with post-hoc Dunn’s multiple comparison was performed to compare pan-tetraspanin with each single tetraspanin detection, p≥0.05 (ns), p<0.05 (*), p<0.01 (**), p<0.001 (***). a.u. = arbitrary unit.

For serum EVs **(Fig. 6b)**, the pan-tetraspanin detection signal of 7 capture bead populations (i.e. CD42a, CD41b, CD31, CD29, CD105, CD40 and HLA-I) was similar to single CD9 detection signal (pan-t VS αCD9, p≥0.05, ns). For the CD40 and HLA-I capture bead populations both CD63 and CD81 single tetraspanin detections were below the isotype control, indicating that CD40^+^ and HLA-I^+^ serum EVs only expressed CD9. For CD42a, CD41b and CD31 capture bead populations, besides CD9, also CD63 single detection signals were measured, while CD81 was not detected. Similarly, serum EVs captured with CD62P showed a comparable profile. Conversely, serum EVs bound to CD105 capture bead expressed, besides CD9, also CD81 but were negative for single CD63 detection. EVs expressing CD29, ROR1, CD24 and HLA-II were detected with all single tetraspanin antibodies. Remarkably, for serum EVs expressing HLA-II, CD8 and CD44 the main contributor to the pan-tetraspanin signal was CD81; for CD44-positive serum EVs, CD81 was the sole contributor to the pan-tetraspanin signal, while for CD8-positive EVs also CD63 was detected. CD133-1-expressing serum EVs were detected with single CD63 and CD81 antibodies, while CD9 was not detectable.

Comparison of the tetraspanin signals on EVs positive for markers detected in both milk and serum EVs (i.e. HLA-I, HLA-II, CD24, CD29, CD44, CD105, CD133-1 and ROR1) unveiled similar tetraspanin signals for EVs expressing HLA-II, CD24, CD29, CD44 and ROR1. For both milk and serum HLA-I EVs, CD63 was not detected and CD9 contributed largely to the pan-tetraspanin signal. However, in contrast to milk EVs, which also expressed CD81, CD81 was not detected on HLA-I^+^ serum EVs. For CD133-1-expressing EVs, in contrast to milk EVs, serum EVs were negative for CD9. Conversely, CD105-expressing milk EVs were negative for CD9, while CD105 serum EVs were negative for CD63 (**Fig. 6a** **and** **6b**).

In conclusion, the comparison of pan-tetraspanin detection with single tetraspanin detection unveiled different tetraspanin profiles depending on the capture antibody and allowed to identify similar and different capture antibody-tetraspanin associations between EVs from milk and serum.

Based on EV proteomics analysis and intracellular trafficking, Kowal *et al.*^11^ and Mathieu *et al.*^12^ recently proposed the use of combinations of CD9, CD63 and/or CD81 for the definition of EV subsets on the basis of their intracellular origin. More specifically, CD9^+^CD63^+^CD81^+^ EVs and CD63^+^CD9^low^ EVs carrying endosomal proteins qualify as exosome (endosomal origin); CD9^+^CD63^−^CD81^−^ and CD9^+^CD81^+^CD63^low^ small EVs mainly bud from the plasma membrane. The subpopulations of CD9 single positive and CD81 single positive small EVs probably form at the plasma membrane and early endocytic locations. Platelet-derived EVs characterized by a CD9^+^CD63^+^CD81^−^ profile are more difficult to classify following the criteria by Kowal *et al.*^11^ and Mathieu *et al.*^12^. However, based on previous observations indicating that platelet-derived EVs in the size range of 30-100nm, expressing CD9, CD63, TSG101, ALIX, CD31, CD41b, CD42a, CD62P and PF4 originate from multi-vesicular bodies^35,36^, we classified platelet-derived serum EVs as endosomal-derived. Based on the cell of origin data as described in **Figure 4** and the tetraspanin distributions described in **Figure 6**, we present a model of the potential cell of origin for the different markers detected on milk EVs and serum EVs connected to proposed EV biogenesis routes^11,12^ **(Fig. 7)**.

**Figure 7.**
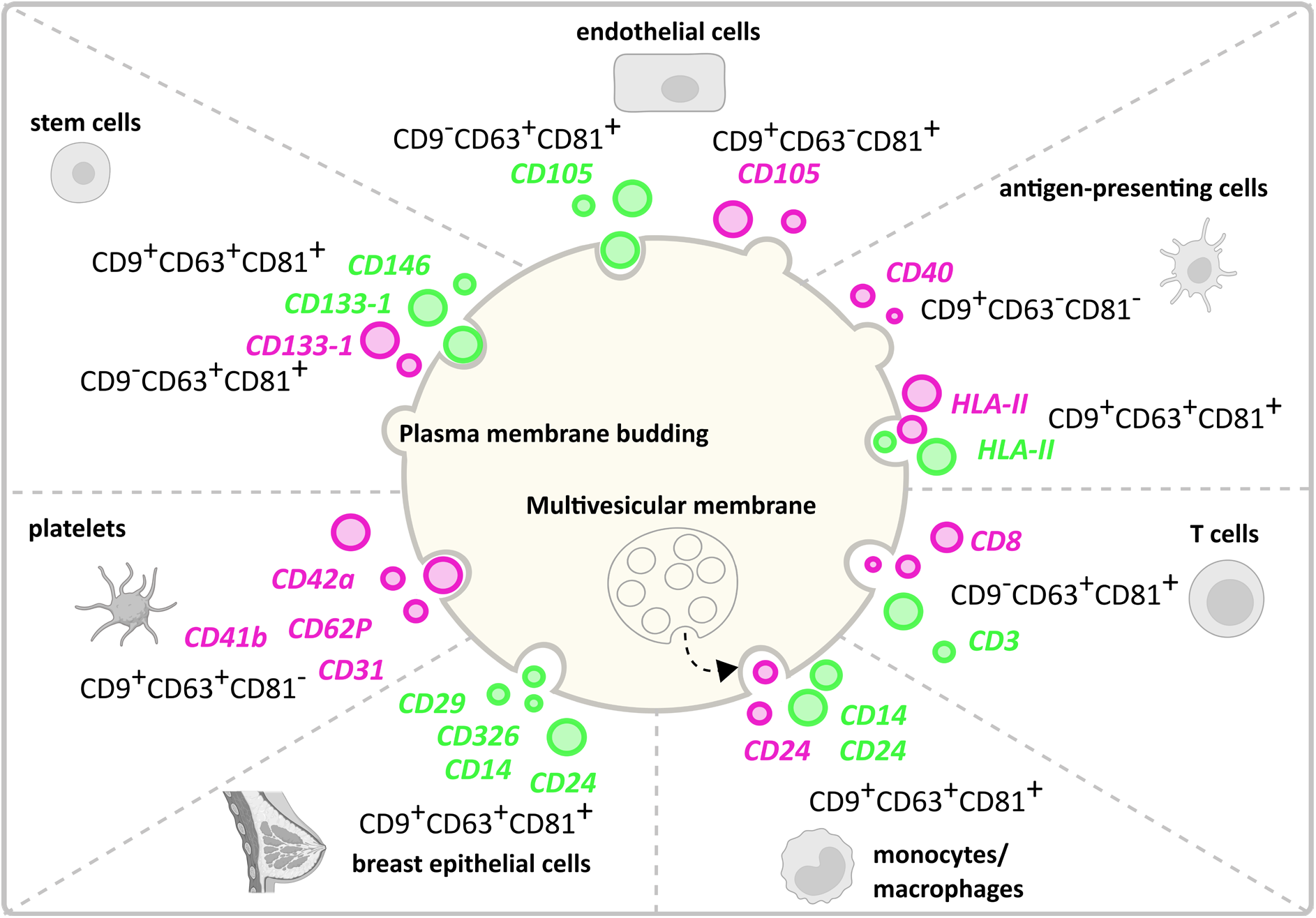
Graphical representation of the potential cell of origin and biogenesis route of milk EVs and serum EVs. This model connects potential cell of origin data described in Figure 4 and the tetraspanin distributions described in Figure 6. Milk EV (green) and serum EV (pink) cell markers are depicted in combination to their association with CD9, CD63 and CD81, from which we propose their biogenesis route based on Kowal *et al.*^11^ and Mathieu *et al.*^12^ study. Created with BioRender.com and Inkscape 1.2.1.

## Materials and methods

### Milk and serum donors

Human milk and serum were collected upon written consent from lactating women who had given birth to a full-term newborn via vaginal delivery. These women were enrolled in the Comparison of Human Milk Extracellular Vesicles in Allergic and Non-allergic Mothers (ACCESS) study (NL 47426.099.14; RTPO 914), approved by the local Medical Ethics Committee (Martini Hospital, Groningen, The Netherlands). Mothers were classified as “allergic” when total serum IgE ≥ 50 kU/L and/or specific IgE was detected for grass pollen, tree pollen, house dust mite, cat dander, or dog dander by positive (≥0.35 kU/L) Phadiatop assay (Thermo Scientific, Uppsala, Sweden). In clinical practice, 0.35 kU/L has commonly been used as a cut off (<0.35 kU/L is considered < limit of detection -LOD-). For this paper, paired milk and serum from n=9 women (n=4 allergic; n=5 non-allergic) were selected based on sample availability. Information on donors is reported in **Supplementary Table S1**.

### Human milk and serum collection

Paired milk and serum samples were collected at the same day during a hospital check-up visit. Milk was collected in sterile Medela’s BPA-free human milk collection containers, as previously described^15^. Shortly, milk was prevented from cooling down and was processed within 20 minutes after collection by 2 rounds of centrifugation at 3,000x*g* for 10 minutes at 23°C (Beckman Coulter Allegra X-12R, Fullerton, CA, USA) to deplete cells and fat. Next, cell and fat-free milk supernatants were stored at −80°C. Peripheral blood (6 ml) was collected in serum-separating tubes. Blood was allowed to clot by leaving it undisturbed for 30 minutes at room temperature. Blood cells and clotted blood were separated from serum by centrifugation at 1,000x*g* for 10 minutes at 23°C and serum was stored at −80°C.

### Preparation of EV samples from human milk and serum

EVs of paired milk and serum samples were isolated using an identical protocol^16^, with minor modifications. In short, equal volumes of stored cell and fat-free milk supernatant and serum (2.5 - 3 ml) were thawed at 37°C, transferred into polyallomer SW60 tubes (Beckman Coulter) and centrifuged at 4°C for 30min at 5,000x*g* and subsequently at 10,000x*g* (Beckman Coulter Optima XPN-80 with a SW60 rotor). Next, 3.5 - 4 ml aliquots of the resulting 10,000x*g* supernatants were added up with 3-2.5 ml 1X PBS (Gibco, Carlsbad, CA, USA) to reach a final volume of 6.5 ml and were transferred on top of a 60% - 10% iodixanol gradient (Progen Biotechnik GmbH, Heidelberg, Germany) in a polyallomer SW40 tube (Beckman Coulter). Gradients were ultracentrifuged at 192,000x*g* for 16 - 18 hours at 4°C (Beckman Coulter Optima XPN-80 with a SW40Ti rotor). Gradient fractions of 500 μl were collected and EV-containing fractions (densities 1.06 - 1.19 g/ml) were pooled. Subsequently, the sample was further purified and Iodixanol was removed by size exclusion chromatography using a 20 ml column (Bio-Rad Laboratories, Hercules, CA, USA) packed with 20 ml Sephadex *g* 100 (Sigma-Aldrich, St. Louis, MO, USA). Fractions of 1 ml were eluted with phenol red free RPMI 1640 medium (Gibco, Carlsbad, CA, USA). EV-containing eluates 3 - 9 (7 ml in total) were pooled, aliquoted and frozen at −80°C until further use.

### Nanoparticle Tracking Analysis (NTA)

Purified milk and serum EV preparations were quantified by NTA using a NanoSight NS300 instrument (Malvern Instruments Ltd, UK) equipped with a 405 nm laser and a high sensitivity sCMOS camera. The NanoSight NTA software version 3.2.16 was used for data acquisition and processing. A NanoSight Syringe Pump (Malvern Instruments Ltd, UK) was used for flow mode measurements, which allows a continuous slow flow of the sample through the measurement chamber. Milk EV and serum EV samples required respectively 1:100 and 1:10-1:50 dilution in endotoxin-free Dulbecco’s 1X PBS without Ca^++^ and Mg^++^ (TMS-012-A Millipore) to reach the recommended concentration of 1*10^8^- 1*10^9^ particles/ml and number of particles in frame between 10 and 110. For each EV sample n=3 60 seconds video captures were acquired with a syringe pump flow rate of 20 AU and controlled temperature set at 20°C. Captures were obtained with camera level 13, camera shutter 1232 and camera gain 219 (default). Capture processing was performed with detection threshold 9, viscosity set to water (0.9cP) and automatic blur and minimum track length. The minimum number of valid tracks per EV sample was set at 1000, as recommended^22^.

### Multiplex bead-based flow cytometric analysis of EV surface proteins by MACSPlex Exosome Kit

Purified milk and serum EVs were subjected to multiplex bead-based analysis by flow cytometry using the human MACSPlex Exosome Kit (Miltenyi Biotec, 130-108-813). EV samples were processed overnight in the MACSPlex Filter Plate as described in the manufacturer’s instructions and in a previously published paper on a systematic evaluation of the assay^19^. Particle counts determined by NTA were used to calculate the input amount of EVs. For the optimization experiment five different input numbers of EVs were chosen in the range of 10^8^ - 10^9^ for milk EVs and 5*10^7^ - 5*10^8^ for serum EVs. All subsequent experiments were performed with 5*10^8^ EVs as input for both milk and serum EVs. To achieve this, the volume of milk and serum EV samples containing the appropriate number of EVs was diluted with MACSPlex buffer to a final volume of 120 μl and added to pre-wet and drained wells of the MACSPlex 96-well 0.22 μm filter plate. 15 μl of MACSPlex Exosome Capture Beads (containing 39 antibody-coated, hard-dyed bead populations) were added to each sample for overnight incubation (17-18 hours) on an orbital shaker at room temperature protected from light. Beads were washed by adding 200 μl of MACSPlex buffer to each sample and wells were drained by centrifuge at 300×*g* for 3 minutes at 23°C. For detection of EVs bound to capture beads, 5 μl of APC-conjugated CD9, CD63 and/or CD81 detection antibodies were added to each well containing 135 μl of MACSPlex buffer and incubated for 1 hour on an orbital shaker at room temperature protected from light. Antibody clones and concentrations used in the kit are proprietary knowledge of Miltenyi Biotec. EV detection was performed with the three detection antibodies used separately (single detection) or together (pan-tetraspanin detection). Samples were washed with 200 μl of MACSPlex buffer and centrifuged at 300×*g* for 3 minutes at 23°C. Next, samples were resuspended in 200 μl of MACSPlex buffer, incubated for 15 minutes on an orbital shaker at room temperature protected from light and centrifuged at 300×*g* for 3 minutes at 23°C. Finally, samples were resuspended in 150 μl of MACSPlex buffer and transferred to a V-bottom 96-well plate for flow cytometric analysis. As negative control RPMI diluted in MACSPlex buffer underwent the same procedure as EV samples to determine non-specific signals. Per well, 135 μl were acquired corresponding to 8,000 - 17,000 recorded beads (**Supplementary Table S2)**. The 39 capture bead populations were distinguished from each others by the 488 nm laser (optical filter 525/40 nm, FITC) and the 561 nm laser (optical filter 585/42 nm, PE), and EVs were detected using the 638 nm laser (optical filter 660/10 nm, APC) of a CytoFLEX LX Flow Cytometer (Beckman Coulter). For data acquisition CytExpert software version 2.1 was used. FlowJo^TM^ software version 10.8.1 (BD Life Sciences) was used for flow cytometry data analysis. Processing of the raw APC-median fluorescence intensity (MFI APC) of all 39 capture bead populations was performed as follows: background correction was performed by subtracting the respective MFI APC values of matched buffer controls, as well as the fluorescent intensity of the corresponding isotype controls (mIgG1, REA). Protein signals with MFI > MFI of isotype controls were considered positive. The other proteins were not used for further analyses. For the generation of part-of-a-whole graphs, to assess the cell origin of EVs, background and isotype subtracted MFI were normalized to 100.

### Statistical analysis

All statistical analyses were performed with GraphPadPrism 9.0.0 (GraphPad Software, San Diego, California USA). Data normality was assessed by Shapiro-Wilk test and d’Agostino-Pearson test. Non-normally distributed variables were expressed as median and analysed by Mann-Whitney test or Kruskal-Wallis test followed by Dunn’s multiple comparison, as appropriate. Paired samples were analysed by Wilcoxon test with FDR. A probability value of <.05 was considered statistically significant.

## Discussion

In this study we used the multiplex bead-based flow cytometry MACSPlex assay to characterize the surface protein profiles of purified EVs, isolated with the same protocol from paired human milk and serum samples. Since 2018, this assay has been used for surface protein profiling of EVs from various body fluids, i.e. blood plasma^19,37–41^, serum^19,42–44^, urine^44^, cerebral spinal fluid^38^ and lymphatic drain fluid^45^ of healthy subjects and patients, using a wide variety of EV enrichment/purification protocols, EV quantification methods and input EV amounts. To our knowledge no studies have been reported on the use of this assay on EVs purified from different body fluids collected on the same day from the same donor, isolated by similar protocols and analysed by pan-tetraspanin as well as single tetrapsanin detection. Analysis of the pan-tetraspanin detection signal (simultaneous detection of CD9, CD63 and CD81) of the CD9, CD63 and CD81 capture bead populations showed that the MFI value for CD9 and CD63 were comparable in milk EVs and serum EVs, while the signal for CD81 capture beads was significantly higher in milk EVs compared to serum EVs. By using the same number of input EVs, the higher CD81 signal on milk EVs could reflect either a higher number of CD81^+^ EVs in milk compared to serum, more CD81 molecules per EV or the presence of EV subsets with high expression levels of CD81. Previously, it has been reported that CD81 was not present on 200,000g/100nm EVs isolated from different cell lines, while it was detectable on 100,000g/200nm EVs, suggesting that CD81 was expressed on larger particles^46^. Accordingly, NTA-based size distribution of EVs used in our study show that milk EVs (234 ± 13 nm) are larger than serum EVs (105 ± 10 nm). Moreover, it was shown by flow cytometry that CD81 MFI signal was 100-1000 times higher in milk EVs compared to EVs purified from plasma^47^, and that CD81^+^ EVs were the rarest EV population in plasma and serum^48^. Furthermore, it has been reported that platelets do not express CD81^49^ and accordingly, platelet-derived EVs were hardly detected with CD81 beads^23^ and were negative in CD81 by westernblotting^50^. As expected, our data show an enrichment of proteins associated with platelets and platelets activation in serum EVs, such as CD41b, CD42a, P-selectin/CD62P and PECAM-1/CD31^50^, an indicator of the presence of platelets-derived EVs. Besides the presence of platelet-derived EVs in serum, we also identified EVs derived from immune cells, albeit their contribution to the total detection signal in the assay was much lower. Also in milk EVs we could detect clear signatures of immune cells-derived EVs, but not blood signatures, thus suggesting that EVs from milk are not likely to come directly from blood.

In line with previously research^51,52^, our data show that milk EVs carry HLA-II, CD14, CD24 and CD3 which reinforce the hypothesis of transfer of immunological features by EVs from mother to infant. In accordance with our previous data^52^, besides immune cells-derived EVs we could detect clear signatures of breast luminal epithelial cells-derived EVs and mesenchymal milk stem cells-derived EVs in milk. Of note, most proteins are not exclusively expressed on one cell type, i.e. CD14 is described as marker of monocytes but is also expressed on mammary epithelial cells^31^. Similarly, CD24 has been associated to both B cells and luminal epithelial cells^32^, and both CD24 and CD326 (EpCAM) are highly expressed on lactocytes^8,28^. Our analysis of milk EVs showed that CD24 and CD326 were two of the highest expressed EV proteins. For defining the potential cell of origin of EVs in milk and serum, we selected the most likely cellular source. However, we cannot exclude that some EVs, expressing markers present on a variety of different cell types, can be derived from other cell types than the ones indicated in our model.

In our study investigating the surface protein profiles of milk and serum EVs from non-allergic (n=5) and allergic (n=4) donors, we observed a tendency for non-allergic milk EVs towards higher MFI compared to allergic EVs, which was not observed in EVs from serum. However, significant differences between clinical groups were not found.

As clearly shown in our current study, besides pan-tetraspanin EV detection, single tetraspanin-detection in relation to specific protein markers can yield important insights on EVs, however this approach is frequently disregarded. By comparing pan-tetraspanin detection (i.e. αCD9, αCD63 and αCD81 antibody cocktail) and single tetraspanin detection (i.e. αCD9, αCD63 or αCD81 single antibody), here we unveiled characteristic tetraspanin profiles for different protein capture bead populations. For example, serum EVs captured with the antigen presenting cell marker HLA-II expressed all three tetraspanins CD9, CD63 and CD81, suggesting that HLA-II^+^ EVs originate and are released via the endosomal route. Conversely, serum EVs expressing the antigen presenting cell associated co-stimulatory protein CD40 were exclusively detected with αCD9 antibody, hinting at their origin from the plasma membrane. These findings suggest the presence of different subsets of antigen presenting cells-derived EVs in serum. On the other side, T cell-associated markers CD3 detected on milk EVs, and CD8 detected on serum EVs were characterized by the same tetraspanin distribution, i.e. positive for CD81 and CD63 but negative for CD9, suggesting that these T cells-derived EVs are from endosomal origin. Finally, milk EVs captured by HLA-I did not express CD63, as opposite to a previous study^53^ showing that CD63-bead bound milk EVs and HLA-I-bead bound milk EVs positively stained for HLA-I and CD63, respectively. This discrepancy can be caused by differences in milk biobanking protocols, e.g. biobanking full milk^53^ versus cell and fat-depleted milk^18^, EV sample preparation, or affinity/avidity for the antibodies used.

In conclusion, the detailed information on specific tetraspanin distribution in conjunction with surface proteins of interest can be exploited for EV-based biomarker profiling in order to unveil and define more robust EV subsets of interest. Furthermore, this knowledge can be used to define and exclude EV subsets in different body fluids that can potentially mask EVs of interest, thus greatly facilitating the analysis of rare EV subsets according to the principle “remove the hay to find the needle”.

## Additional information

## Acknowledgements

We thank Ger Arkesteijn for his constant support and the use of the Flow Cytometry Facility (Faculty of Veterinary Medicine, Utrecht University, The Netherlands) and Johan Garssen (Department of Pharmaceutical Sciences, Faculty of Science, Utrecht University, The Netherlands and Nutricia Research Centre for Specialized Nutrition, The Netherlands), Frank A. Redegeld (Department of Pharmaceutical Sciences, Faculty of Science, Utrecht University, The Netherlands), Ruurd H. van Elburg (Department of Pediatrics, Emma Children’s Hospital/Academic Medical Center, Amsterdam, The Netherlands), and Arianne van Bruggen-de Haan (Paediatric Allergy Centre, Department of Paediatrics, Martini Hospital, Groningen, The Netherlands) for their contributions to the clinical ACCESS study from which biobanked samples were used in this study.

## Author contributions

A.G. designed and performed experiments, analysed data and wrote the manuscript. M.H.M.W. supervised the research, designed the clinical study and experiments and wrote the manuscript. G.N.M. designed the clinical study and was involved in subject recruitment. M.J.C.H designed and performed milk sample preparation for biobanking and revised the manuscript. All authors critically reviewed and edited the manuscript.

## Data availability

Raw multiplex data are shared in Supplementary Tables. Additional information for reanalysis may be requested from the lead contact. We have submitted all relevant data of our experiments to the EV-TRACK knowledgebase^54^ (EV-TRACK ID: EV220315).

## Supplementary Information

Additional supporting information can be found online in the corresponding section at the end of the article.

Figures: S1-S6

Tables: S1-S4

