## Supplementary information for "Surface protein profiling of milk and serum extracellular vesicles unveil body fluid and cell-type signatures and insights on vesicle biogenesis"

3 TRAIN-EV Marie Skłodowska-Curie Action-ITN

*****

### SUPPLEMENTARY INFORMATION

| Supplementary Table S1. Clinical parameters of milk and serum donors | | | | | | | | | |
|---|---|---|---|---|---|---|---|---|---|
| ***Donor number*** | ***1*** | ***2*** | ***3*** | ***4*** | ***5*** | ***6*** | ***7*** | ***8*** | ***9*** |
| ***Moternal age*** *(years)* | 30 | 34 | 34 | 31 | 32 | 36 | 25 | 31 | 33 |
| ***Lactational stage*** *(weeks post-partum)* | 9 | 5 | 6 | 5 | 6 | 8 | 7 | 5 | 6 |
| ***Number of full term deliveries*** | 2 | 4 | 2 | 2 | 2 | 1 | 1 | 1 | 3 |
| ***Health status*** | A | NA | A | NA | A | NA | A | NA | NA |
| ***Total IgE*** *(kU/L)* | 73.6 | 24.7 | 243 | 27.8 | 141 | 38.9 | 81.9 | 30.4 | 9.6 |
| ***grass pollen IgE*** *(kU/L)* | < LOD | < LOD | 0.97 | < LOD | < LOD | < LOD | 1.3 | < LOD | < LOD |
| ***tree pollen IgE*** *(kU/L)* | 7.6 | < LOD | 40 | < LOD | < LOD | < LOD | 7.7 | < LOD | < LOD |
| ***house dust mite IgE*** *(kU/L)* | 1.0 | < LOD | 4.2 | < LOD | 3.8 | < LOD | 4.4 | < LOD | < LOD |
| ***cat dander IgE*** *(kU/L)* | < LOD | < LOD | < LOD | < LOD | < LOD | < LOD | 0.49 | < LOD | < LOD |
| ***dog dander IgE*** *(kU/L)* | < LOD | < LOD | < LOD | < LOD | < LOD | < LOD | 1.1 | < LOD | < LOD |

LOD= limit of detection
A= allergic
NA=non-allergic

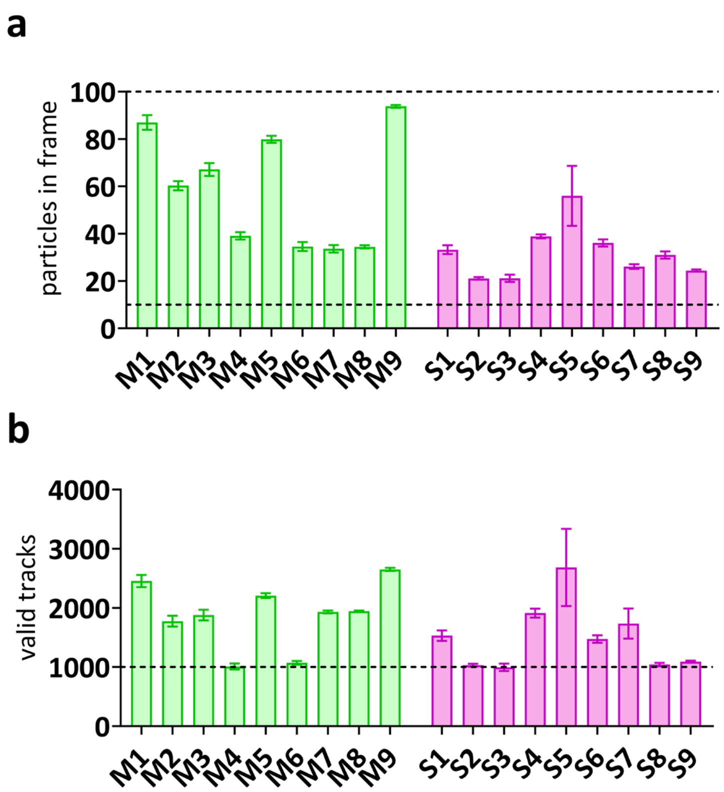


**Supplementary Figure S1. Technical quality controls for NTA.** To ensure reliable measurements, milk and serum EVs were diluted to reach 10-110 particles in frame **(a)** and at least 1,000 valid tracks as average of 3 captures per EV sample (**b**). The results shown are derived from n=9 individual milk samples (M1-M9) and paired serum samples (S1-S9).

#### Supplementary Table S2. Raw bead counts and percentages of capture signals

| samples | acquired events | beads | single beads | APC+ beads (FITC) | APC+ beads (PE) | APC+ beads (FITC) | APC+ beads (PE) |
|---|---|---|---|---|---|---|---|
| | (counts) | (counts) | (counts) | (counts) | | (% of single beads) | |
| *buffer* | 17256 | 15947 | 15929 | 196 | 195 | 1.23 | 1.22 |
| *$10^9$ milk 1* | 15579 | 11302 | 11077 | 3415 | 3412 | 30.8 | 30.8 |
| *$10^9$ milk 2* | 17325 | 13074 | 13007 | 4074 | 4065 | 31.3 | 31.3 |
| *$10^9$ milk 3* | 18106 | 13349 | 13290 | 4527 | 4522 | 34.1 | 34 |
| *7.5*$10^8$ milk 1* | 16533 | 12543 | 12415 | 3344 | 3333 | 26.9 | 26.8 |
| *7.5*$10^8$ milk 2* | 18137 | 14292 | 14240 | 4129 | 4118 | 29 | 28.9 |
| *7.5*$10^8$ milk 3* | 17686 | 13392 | 13273 | 4198 | 4189 | 31.6 | 31.6 |
| *5*$10^8$ milk 1* | 19146 | 14980 | 14805 | 3508 | 3494 | 23.7 | 23.6 |
| *5*$10^8$ milk 2* | 17308 | 13267 | 13227 | 3359 | 3351 | 25.4 | 25.3 |
| *5*$10^8$ milk 3* | 21644 | 17311 | 17156 | 5055 | 5049 | 29.5 | 29.4 |
| *2.5*$10^8$ milk 1* | 12010 | 8988 | 8884 | 1925 | 1926 | 21.7 | 21.7 |
| *2.5*$10^8$ milk 2* | 11750 | 8779 | 8764 | 2040 | 2037 | 23.3 | 23.2 |
| *2.5*$10^8$ milk 3* | 10885 | 7913 | 7790 | 2119 | 2117 | 27.2 | 27.2 |
| *$10^8$ milk 1* | 14984 | 11252 | 11196 | 2286 | 2278 | 20.4 | 20.3 |
| *$10^8$ milk 2* | 16818 | 12852 | 12792 | 2629 | 2621 | 20.6 | 20.5 |
| *$10^8$ milk 3* | 16757 | 13063 | 13010 | 2955 | 2948 | 22.7 | 22.7 |
| *5*$10^8$ serum 1* | 18031 | 12344 | 12000 | 3784 | 3766 | 31.5 | 31.4 |
| *5*$10^8$ serum 2* | 14758 | 10369 | 10316 | 4004 | 3992 | 38.8 | 38.7 |
| *5*$10^8$ serum 3* | 19132 | 13693 | 13530 | 4035 | 4023 | 29.8 | 29.7 |
| *2.5*$10^8$ serum 1* | 17481 | 12305 | 12041 | 2856 | 2845 | 23.7 | 23.6 |
| *2.5*$10^8$ serum 2* | 15387 | 11029 | 10974 | 3400 | 3379 | 31 | 30.8 |
| *2.5*$10^8$ serum 3* | 16943 | 12510 | 12120 | 2928 | 2925 | 24.2 | 24.1 |
| *$10^8$ serum 1* | 17796 | 13336 | 13230 | 2291 | 2280 | 17.3 | 17.2 |
| *$10^8$ serum 2* | 15949 | 11959 | 11711 | 2584 | 2569 | 22.1 | 21.9 |
| *$10^8$ serum 3* | 13313 | 9847 | 9512 | 1972 | 1966 | 20.7 | 20.7 |
| *7.5*$10^7$ serum 1* | 15056 | 10801 | 10566 | 1566 | 1557 | 14.8 | 14.7 |
| *7.5*$10^7$ serum 2* | 13441 | 9759 | 9590 | 1859 | 1846 | 19.4 | 19.2 |
| *7.5*$10^7$ serum 3* | 11509 | 8257 | 8053 | 1552 | 1549 | 19.3 | 19.2 |
| *5*$10^7$ serum 1* | 18373 | 13975 | 13843 | 1722 | 1710 | 12.4 | 12.4 |
| *5*$10^7$ serum 2* | 18019 | 14171 | 14001 | 1638 | 1618 | 11.7 | 11.6 |
| *5*$10^7$ serum 3* | 15502 | 12136 | 11801 | 1875 | 1868 | 15.9 | 15.8 |

**Supplementary Table S2. Raw bead counts and percentages of MACSPlexbead capture signals.** Acquired events is the number of events recorded in 135 µl of volume at medium flow rate (30 sec/min). Beads were gated and single beads were the events used as input for the analysis of count or relative percentage of $APC^+$ bead populations (EVs bound to beads). Beads are distinguished from each others by their respective fluorescence characteristics using the lasers and optical filters for FITC and PE. Data were obtained from the gating strategy in Supplementary Fig. S2a. Data are shown in the bar graphs of Supplementary Fig. S2b.

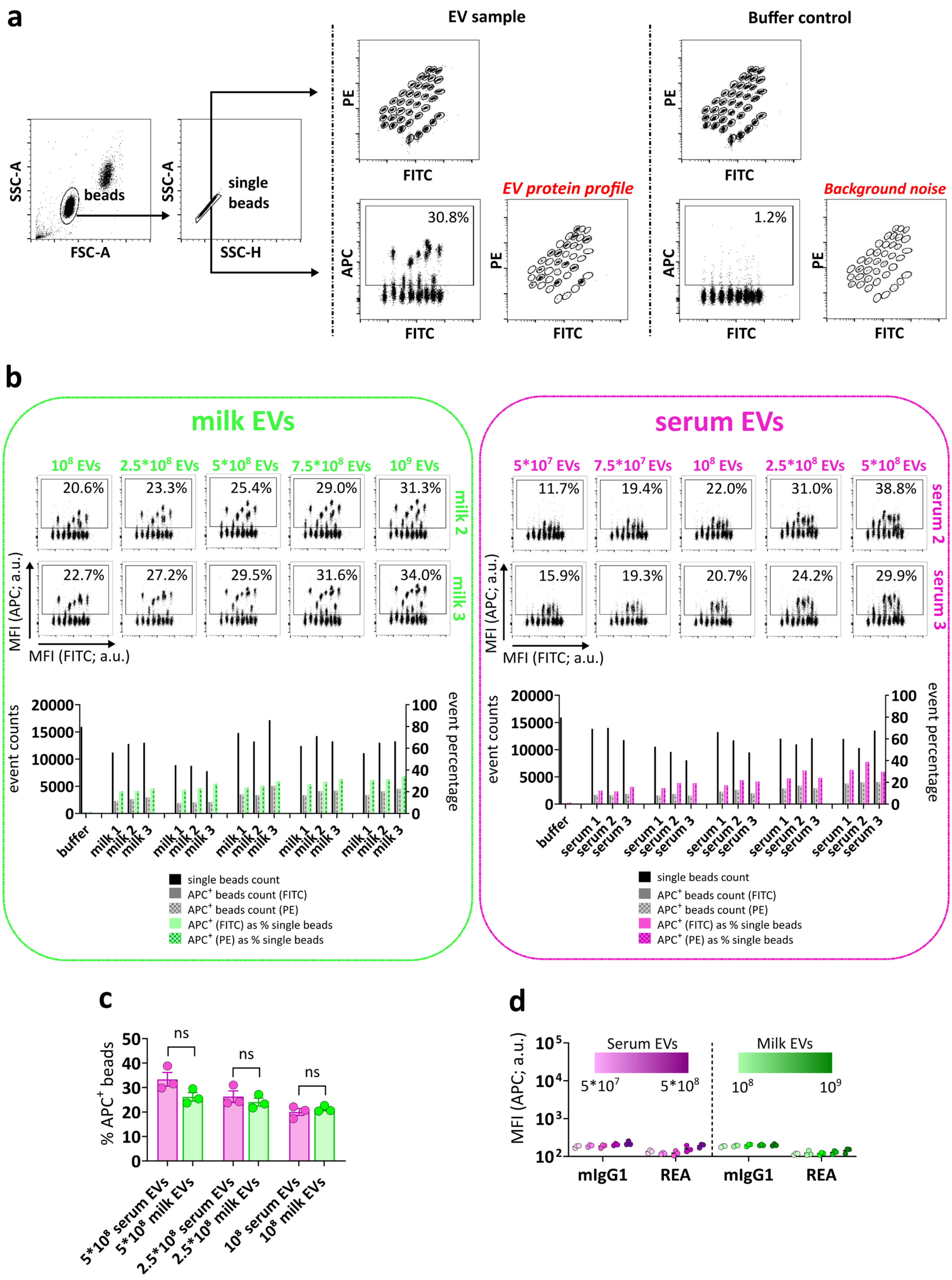
a
EV sample
Buffer control
SSC-A
FSC-A
beads
SSC-A
SSC-H
single beads
PE
FITC
APC
30.8%
EV protein profile
1.2%
Background noise
b
milk EVs
serum EVs
MFI (APC; a.u.)
MFI (FITC; a.u.)
milk 2
milk 3
serum 2
serum 3
event counts
event percentage
single beads count
APC+ beads count (FITC)
APC+ beads count (PE)
APC+ (FITC) as % single beads
APC+ (PE) as % single beads
c
% APC+ beads
ns
5*10^8 serum EVs
5*10^8 milk EVs
2.5*10^8 serum EVs
2.5*10^8 milk EVs
10^8 serum EVs
10^8 milk EVs
d
MFI (APC; a.u.)
Serum EVs
Milk EVs
mIgG1
REA

**Supplementary Figure S2. Titration of APC detection signals is proportional to input EV numbers. (a)** Overview of the MACSPlex gating strategies: events in 135ul were recorded at medium flow rate (30sec/min). Beads were gated and single beads were the events used as input for the analysis of $APC^+$ bead populations (EVs bound to beads). Beads are distinguished from each others by their respective fluorescence characteristics using the lasers and optical filters for FITC and PE. **(b)** Flow cytometry dot plots of five input amounts of milk EVs (green) and serum EVs (pink) from donor 2 and 3 depicting the percentage of $APC^+$ beads (relative to single beads). The bar graphs show the counts of single beads and counts and percentages of $APC^+$ beads of three donors (donor 1, donor 2, donor 3). See Supplementary Table S2 for further details. **(c)** Technical control: percentage of $APC^+$ beads (EVs bound to beads) in relation to input number of milk EVs and serum EVs. The graph depicts median with dots of donor 1, donor 2, donor 3) **(d)** Technical control: MFI APC relative to internal isotype controls mIgG1 and REA capture beads at different EV doses. The graph depicts median + min and max values (n=3 donors). a.u. = arbitrary unit.

#### Supplementary Table S3: Raw MFI APC signal (pan-tetraspanin) of all 39 beads

| | *buffer* | *M1* | *M2* | *M3* | *M4* | *M5* | *M6* | *M7* | *M8* | *M9* | *S1* | *S2* | *S3* | *S4* | *S5* | *S6* | *S7* | *S8* | *S9* |
|---|---|---|---|---|---|---|---|---|---|---|---|---|---|---|---|---|---|---|---|
| CD1c | 125 | 125 | 195 | 291 | 213 | 305 | 195 | 245 | 172 | 220 | 222 | 1640 | 271 | 959 | 538 | 228 | 327 | 233 | 322 |
| CD2 | 141 | 204 | 258 | 428 | 297 | 193 | 485 | 314 | 310 | 243 | 202 | 267 | 182 | 209 | 239 | 254 | 265 | 226 | 228 |
| CD3 | 432 | 820 | 1918 | 1804 | 2089 | 2403 | 680 | 2023 | 1055 | 1399 | 295 | 479 | 316 | 228 | 314 | 258 | 248 | 191 | 274 |
| CD4 | 125 | 125 | 171 | 241 | 169 | 116 | 222 | 172 | 230 | 151 | 176 | 152 | 149 | 149 | 189 | 176 | 169 | 165 | 180 |
| CD8 | 101 | 204 | 308 | 422 | 250 | 211 | 444 | 282 | 261 | 252 | 1432 | 2314 | 1491 | 1333 | 1281 | 1766 | 1097 | 1618 | 1523 |
| CD9 | 125 | 6762 | 6639 | 14726 | 7534 | 10658 | 26482 | 8171 | 33644 | 8019 | 5754 | 7724 | 13842 | 2740 | 9595 | 6762 | 14871 | 3230 | 14969 |
| CD11c | 138 | 149 | 132 | 250 | 149 | 187 | 151 | 178 | 176 | 141 | 853 | 263 | 231 | 140 | 176 | 140 | 169 | 165 | 140 |
| CD14 | 305 | 24353 | 25099 | 54009 | 37119 | 33305 | 38141 | 32746 | 126955 | 15878 | 485 | 639 | 687 | 511 | 698 | 399 | 538 | 553 | 493 |
| CD19 | 143 | 163 | 200 | 217 | 217 | 333 | 198 | 196 | 178 | 136 | 280 | 513 | 274 | 259 | 728 | 198 | 261 | 352 | 263 |
| CD20 | 280 | 343 | 373 | 493 | 446 | 570 | 446 | 434 | 483 | 348 | 350 | 731 | 375 | 348 | 738 | 327 | 389 | 432 | 316 |
| CD24 | 162 | 16735 | 6659 | 41808 | 11510 | 29196 | 26042 | 17013 | 40547 | 14207 | 1012 | 970 | 522 | 905 | 3167 | 790 | 978 | 950 | 1257 |
| CD25 | 401 | 195 | 219 | 356 | 346 | 547 | 219 | 288 | 254 | 196 | 273 | 651 | 263 | 348 | 858 | 195 | 274 | 341 | 224 |
| CD29 | 116 | 967 | 918 | 3028 | 1287 | 2651 | 6245 | 1020 | 4442 | 1507 | 2786 | 6700 | 6439 | 1091 | 7772 | 4058 | 6170 | 1115 | 4390 |
| CD31 | 172 | 172 | 198 | 387 | 195 | 329 | 436 | 436 | 442 | 176 | 1052 | 761 | 1693 | 538 | 4574 | 1257 | 1819 | 479 | 2175 |
| CD40 | 256 | 267 | 284 | 493 | 274 | 339 | 717 | 335 | 405 | 288 | 937 | 992 | 2163 | 409 | 4301 | 850 | 1322 | 507 | 2615 |
| CD41b | 337 | 373 | 362 | 517 | 360 | 678 | 474 | 440 | 553 | 360 | 8755 | 6226 | 6558 | 2163 | 14823 | 12356 | 15878 | 3690 | 19741 |
| CD42a | 118 | 156 | 101 | 147 | 140 | 198 | 105 | 132 | 151 | 138 | 8978 | 5806 | 23008 | 1766 | 23790 | 12080 | 26042 | 2320 | 27204 |
| CD44 | 258 | 373 | 339 | 557 | 401 | 466 | 1106 | 448 | 604 | 358 | 393 | 578 | 540 | 483 | 2538 | 735 | 1094 | 528 | 761 |
| CD45 | 269 | 282 | 293 | 483 | 312 | 352 | 430 | 368 | 430 | 288 | 411 | 362 | 483 | 401 | 1644 | 551 | 604 | 345 | 931 |
| CD49e | 215 | 193 | 184 | 265 | 213 | 182 | 200 | 213 | 196 | 228 | 261 | 211 | 219 | 217 | 750 | 207 | 335 | 200 | 356 |
| CD56 | 671 | 815 | 626 | 664 | 566 | 800 | 624 | 825 | 666 | 559 | 428 | 507 | 454 | 432 | 364 | 393 | 354 | 322 | 430 |
| CD62P | 97.7 | 202 | 81.4 | 95.9 | 81.4 | 134 | 85 | 92.3 | 120 | 86.8 | 7126 | 5949 | 17525 | 964 | 15723 | 11111 | 24682 | 3765 | 21667 |
| CD63 | 230 | 6867 | 6931 | 22179 | 9504 | 11147 | 11510 | 6095 | 36619 | 7039 | 10225 | 17525 | 25954 | 3341 | 19417 | 10899 | 13797 | 6598 | 22105 |
| CD69 | 184 | 299 | 289 | 576 | 545 | 1026 | 312 | 379 | 639 | 239 | 479 | 1523 | 553 | 327 | 1287 | 325 | 540 | 572 | 481 |
| CD81 | 191 | 15569 | 20681 | 33758 | 22253 | 25695 | 55509 | 21098 | 64997 | 19035 | 9595 | 8069 | 9223 | 6303 | 10624 | 8171 | 9841 | 9092 | 10726 |
| CD86 | 209 | 265 | 312 | 474 | 379 | 624 | 341 | 329 | 343 | 245 | 243 | 524 | 271 | 269 | 460 | 243 | 295 | 671 | 245 |
| CD105 | 196 | 452 | 356 | 810 | 515 | 778 | 622 | 430 | 2371 | 368 | 1067 | 669 | 669 | 553 | 717 | 485 | 628 | 454 | 530 |
| CD133/1 | 239 | 30197 | 10323 | 43701 | 13355 | 12119 | 29000 | 5516 | 180227 | 17994 | 387 | 1184 | 680 | 515 | 918 | 549 | 648 | 581 | 476 |
| CD142 | 274 | 346 | 337 | 454 | 373 | 389 | 442 | 391 | 495 | 325 | 265 | 314 | 360 | 291 | 324 | 293 | 295 | 335 | 284 |
| CD146 | 235 | 918 | 983 | 1519 | 771 | 1593 | 3659 | 1165 | 1954 | 931 | 609 | 440 | 515 | 393 | 381 | 391 | 327 | 375 | 345 |
| CD209 | 94.1 | 94.1 | 138 | 154 | 101 | 123 | 118 | 136 | 132 | 120 | 112 | 187 | 114 | 95.9 | 158 | 77.8 | 95.9 | 174 | 120 |
| CD326 | 585 | 49245 | 62803 | 86494 | 44452 | 65897 | 161834 | 54195 | 109019 | 56857 | 511 | 3909 | 921 | 622 | 781 | 491 | 517 | 873 | 505 |
| mIgG1 | 202 | 191 | 222 | 370 | 220 | 306 | 362 | 377 | 358 | 217 | 191 | 256 | 200 | 312 | 413 | 295 | 343 | 333 | 202 |
| REA | 132 | 149 | 147 | 158 | 136 | 180 | 101 | 149 | 180 | 143 | 182 | 226 | 189 | 125 | 191 | 118 | 123 | 136 | 143 |
| HLA-I | 462 | 850 | 738 | 1539 | 611 | 581 | 4494 | 637 | 1109 | 536 | 992 | 902 | 1134 | 589 | 2961 | 1555 | 3456 | 553 | 2470 |
| HLA-II | 171 | 16516 | 21885 | 54942 | 30710 | 9873 | 127396 | 12003 | 87999 | 16035 | 1653 | 3114 | 2637 | 3062 | 2234 | 2175 | 3322 | 2192 | 3131 |
| MCSP | 202 | 206 | 248 | 297 | 276 | 385 | 278 | 267 | 252 | 217 | 233 | 520 | 289 | 250 | 377 | 246 | 278 | 346 | 256 |
| ROR1 | 213 | 915 | 1503 | 4058 | 1893 | 2817 | 16194 | 1200 | 3408 | 2371 | 1274 | 3071 | 745 | 820 | 1734 | 405 | 764 | 1124 | 1152 |
| SSEA-4 | 368 | 424 | 397 | 460 | 458 | 752 | 356 | 297 | 470 | 413 | 301 | 764 | 411 | 284 | 530 | 258 | 288 | 310 | 446 |

**Supplementary Table S3. Raw MFI APC signal (pan-tetraspanin) of all 39 bead populations of the MACSPlex Exosome kit.** Raw MFI APC intensities of each capture bead population without background and isotype corrections are indicated. M1-M9 refers to milk EV samples of n=9 individual donors, while S1-S9 are the paired serum EV samples.

#### Supplementary Table S4_Dunn's multiple comparisons test isotype VS proteins

**milk**

| | Mean rank diff. | Adj p value | Significant |
|---|---|---|---|
| CD1c | -62 | >0.99 | ns |
| CD2 | -107 | 0.91 | ns |
| **CD3** | **-198** | **0.001** | ** |
| CD4 | -36 | >0.99 | ns |
| CD8 | -126 | 0.29 | ns |
| **CD9** | **-246** | **<0.001** | *** |
| CD11c | -9.9 | >0.99 | ns |
| **CD14** | **-276** | **<0.001** | *** |
| CD19 | -50 | >0.99 | ns |
| CD20 | -107 | 0.88 | ns |
| **CD24** | **-260** | **<0.001** | *** |
| CD25 | -12 | >0.99 | ns |
| **CD29** | **-213** | **<0.001** | *** |
| CD31 | -93 | >0.99 | ns |
| CD40 | -59 | >0.99 | ns |
| CD41b | -124 | 0.32 | ns |
| CD42a | -51 | >0.99 | ns |
| CD44 | -144 | 0.09 | ns |
| CD45 | -56 | >0.99 | ns |
| CD49e | 0 | >0.99 | ns |
| CD56 | -50 | >0.99 | ns |
| CD62P | -25 | >0.99 | ns |
| **CD63** | **-245** | **<0.001** | *** |
| CD69 | -135 | 0.17 | ns |
| **CD81** | **-269** | **<0.001** | *** |
| CD86 | -92 | >0.99 | ns |
| **CD105** | **-174** | **0.009** | ** |
| **CD133-1** | **-261** | **<0.001** | *** |
| CD142 | -88 | >0.99 | ns |
| **CD146** | **-203** | **<0.001** | *** |
| CD209 | -25 | >0.99 | ns |
| **CD326** | **-291** | **<0.001** | *** |
| **HLA-I** | **-172** | **0.01** | * |
| **HLA-II** | **-269** | **<0.001** | *** |
| MCSP | -46 | >0.99 | ns |
| **ROR1** | **-218** | **<0.001** | *** |
| SSEA-4 | -101 | >0.99 | ns |

**serum**

| | Mean rank diff. | Adj p value | Significant |
|---|---|---|---|
| CD1c | -113 | 0.62 | ns |
| CD2 | -45 | >0.99 | ns |
| CD3 | 0 | >0.99 | ns |
| CD4 | -26 | >0.99 | ns |
| **CD8** | **-210** | **<0.001** | *** |
| **CD9** | **-264** | **<0.001** | *** |
| CD11c | -47 | >0.99 | ns |
| CD14 | -115 | 0.58 | ns |
| CD19 | -88 | >0.99 | ns |
| CD20 | -69 | >0.99 | ns |
| **CD24** | **-189** | **0.002** | ** |
| CD25 | -29 | >0.99 | ns |
| **CD29** | **-240** | **<0.001** | *** |
| **CD31** | **-194** | **0.002** | ** |
| **CD40** | **-180** | **0.005** | ** |
| **CD41b** | **-265** | **<0.001** | *** |
| **CD42a** | **-271** | **<0.001** | *** |
| **CD44** | **-152** | **<0.05** | * |
| CD45 | -114 | 0.6 | ns |
| CD49e | -45 | >0.99 | ns |
| CD56 | 0 | >0.99 | ns |
| **CD62P** | **-267** | **<0.001** | *** |
| **CD63** | **-278** | **<0.001** | *** |
| CD69 | -143 | 0.09 | ns |
| **CD81** | **-270** | **<0.001** | *** |
| CD86 | -67 | >0.99 | ns |
| **CD105** | **-152** | **<0.05** | * |
| **CD133-1** | **-147** | **<0.05** | * |
| CD142 | -23 | >0.99 | ns |
| CD146 | -83 | >0.99 | ns |
| CD209 | -29 | >0.99 | ns |
| CD326 | -58 | >0.99 | ns |
| **HLA-I** | **-179** | **0.006** | ** |
| **HLA-II** | **-231** | **<0.001** | *** |
| MCSP | -50 | >0.99 | ns |
| **ROR1** | **-184** | **0.004** | ** |
| SSEA-4 | -38 | >0.99 | ns |

**Supplementary Table S4. Dunn's multiple comparison test isotype VS protein signals.** Statistical analysis to define proteins below the isotype threshold (non-detected) and above the isotype threshold (expressed on EVs). Significance: ns ($p \geq 0.05$), * ($p<0.05$), ** ($p<0.01$), *** ($p<0.001$).

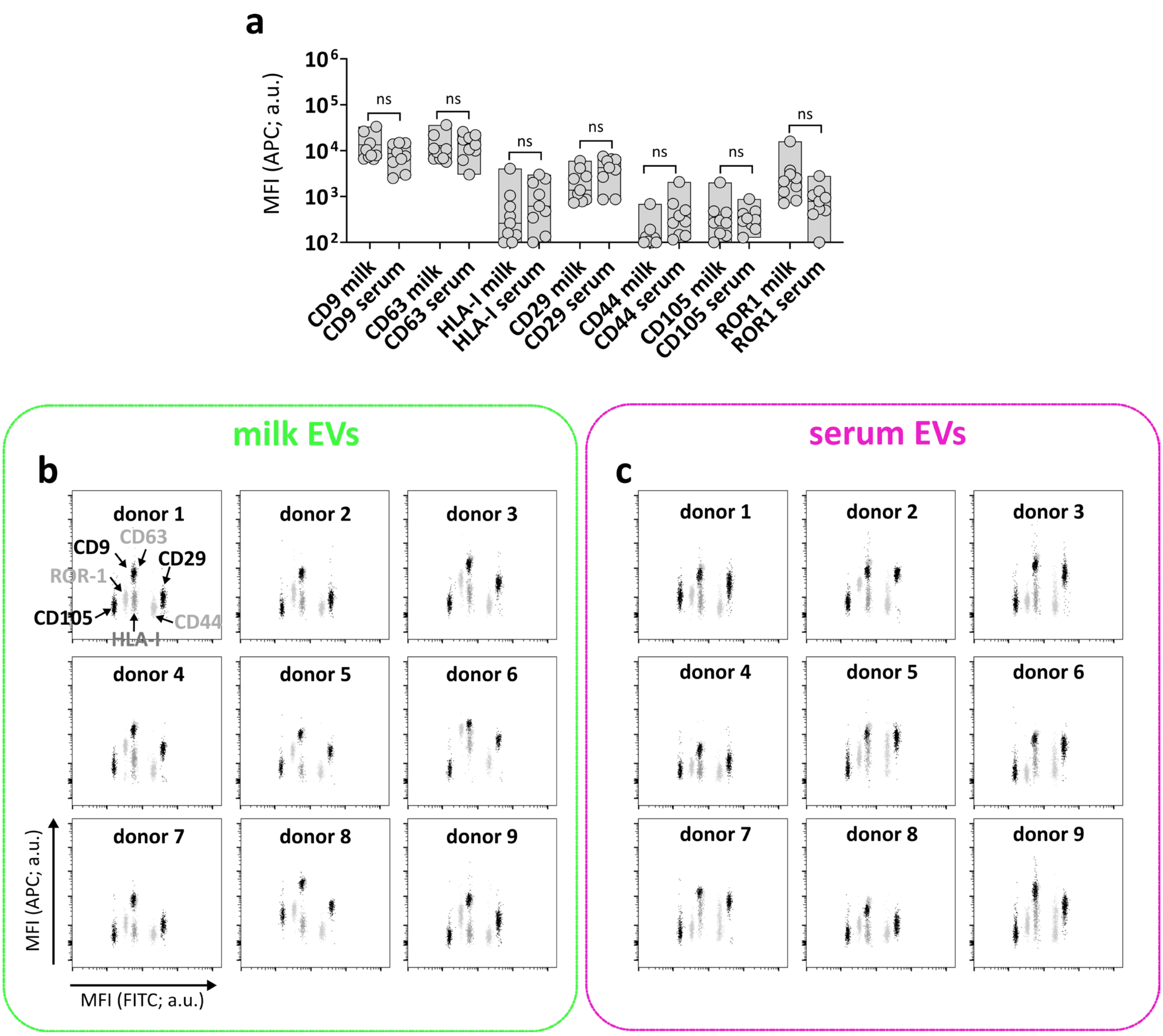


**Supplementary Figure S3. Common milk EV and serum EV proteins with non-significant differences in pan-tetraspanin detection. (a)** Indicated are the MFI APC pan-tetraspanin signals of n=9 individual EV samples per group. **(b)** Corresponding flow cytometry dotplots of the n=9 individual EV samples of proteins depicted in (a). p≥0.05 (ns), non-parametric paired T test (Wilcoxon test). a.u. = arbitrary unit.

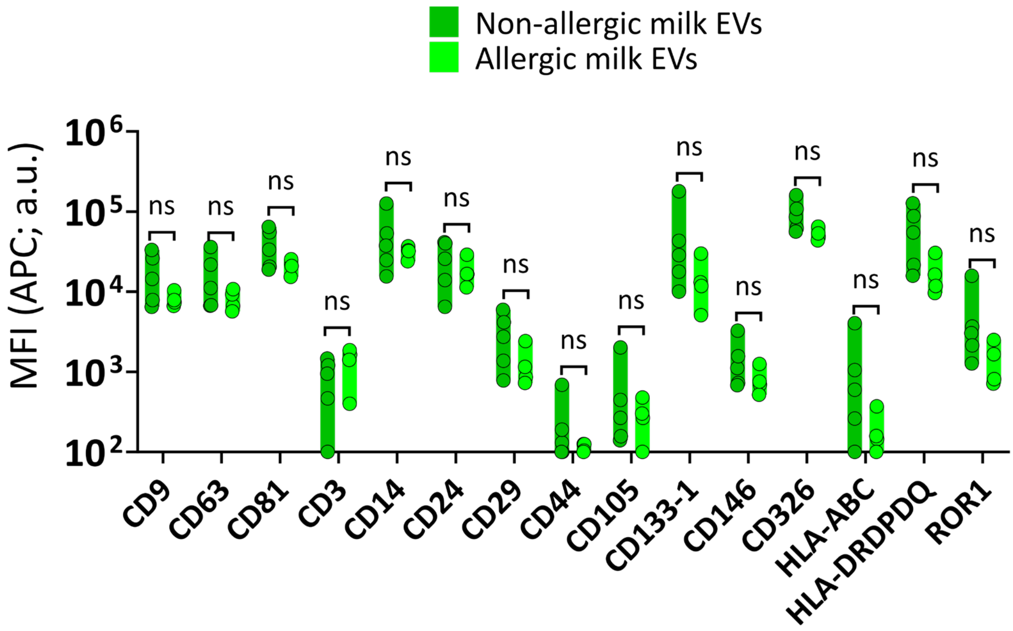


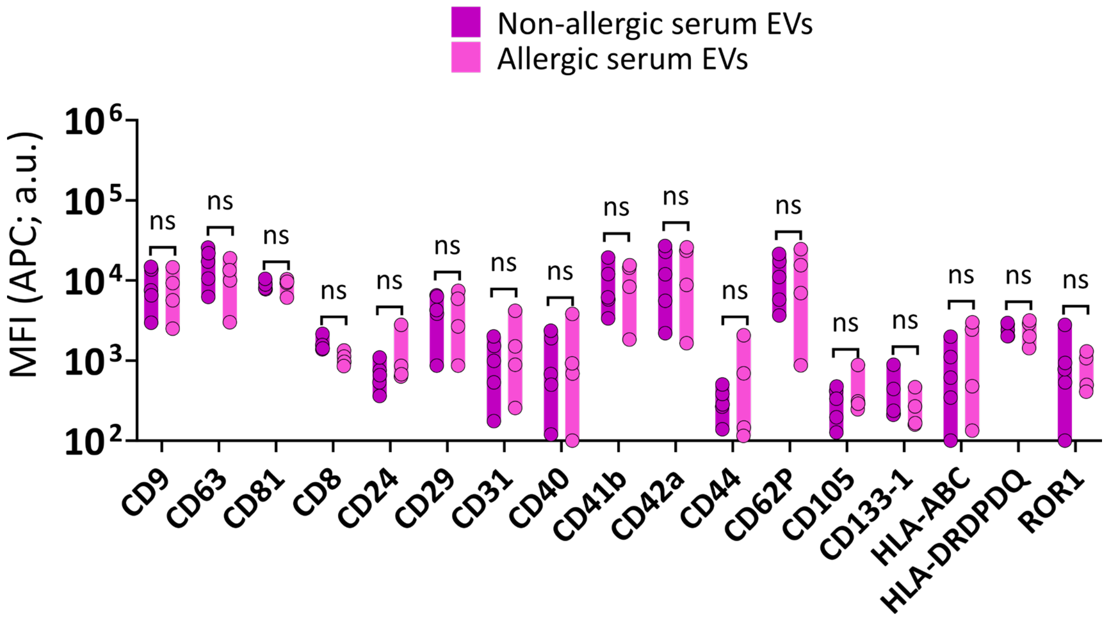


**Supplementary Figure S4. Comparison of EVs from non-allergic and allergic donors.** MFI APC of capture bead populations higher than the respective isotype controls (Figure 3) was compared between milk EVs and serum EVs from non-allergic (n=5 donors) and allergic (n=4 donors) mothers. Significance differences between non-allergic and allergic groups was assessed by multiple non-parametric Mann Whitney test with Holm-Šídák multiple comparison method. a.u. = arbitrary unit.

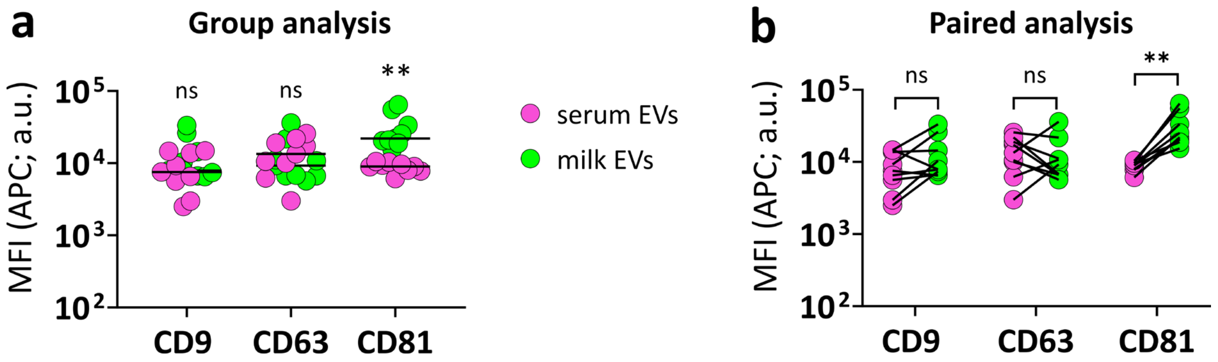


**Supplementary Figure S5. Comparison of CD9, CD63 and CD81 bead capture signals in milk and serum EVs with pan-tetraspanin detection. (a)** Comparison between milk and serum EVs (n=9 samples/group). Lines represent the medians. **(b)** Paired analysis of milk and serum EVs of individual donors (n=9 donors). p≥0.05 (ns), p<0.01 (**), Paired T test nonparametric (Wilcoxon test). a.u. = arbitrary unit.

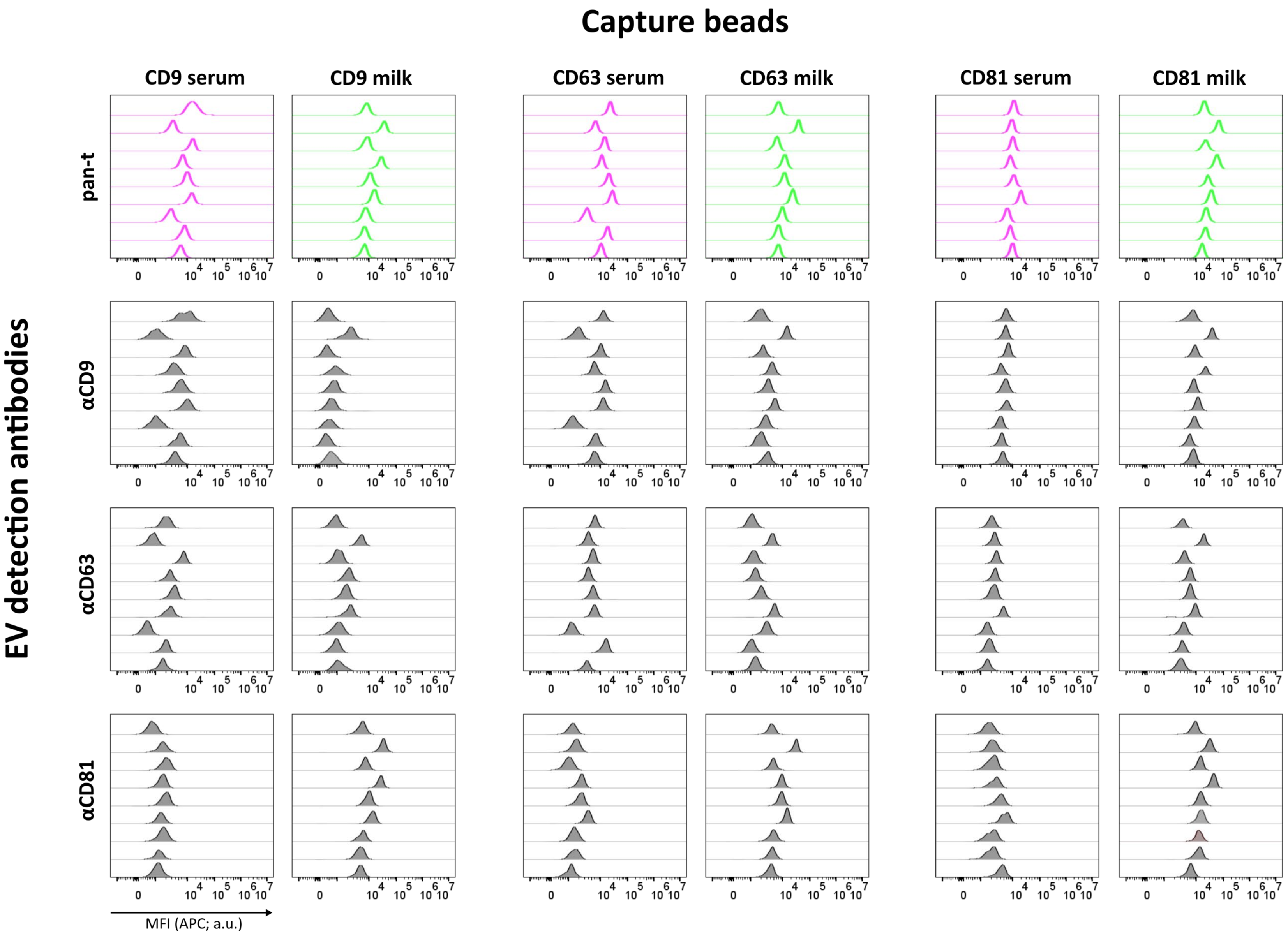


**Supplementary Figure S6.** Flow cytometry histograms representing MFI APC (x-axis) of CD9, CD63, CD81 capture bead populations in milk EVs and serum EVs (n= 9 donors) with pan-tetraspanin detection (pan-t) in comparison to single tetraspanin detection (αCD9, αCD63, αCD81). a.u. = arbitrary unit.

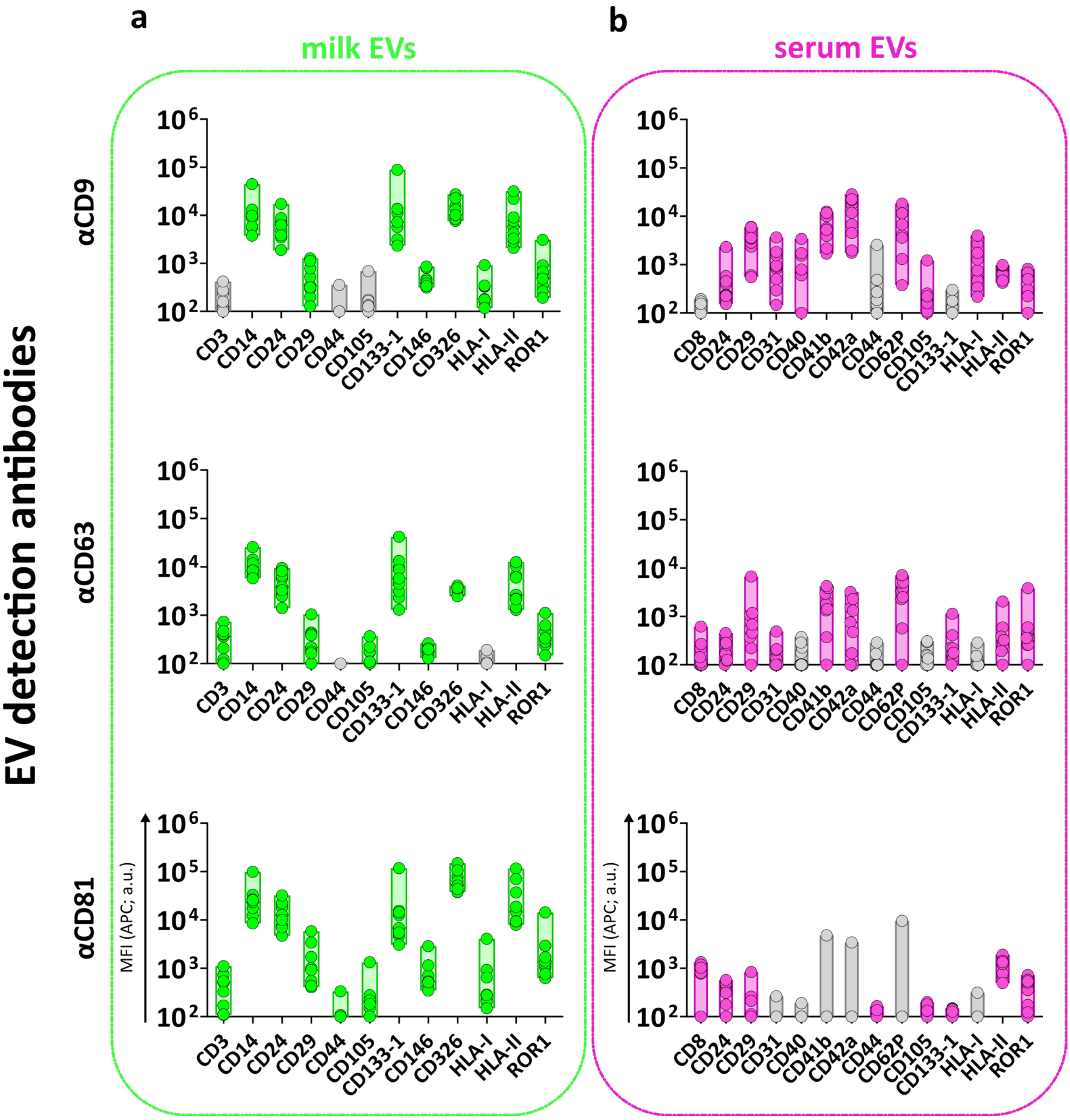


**Supplementary Figure S7. The MFI APC of single tetraspanin detection on EVs bound to capture bead populations identified by pan-tetraspanin detection.** Single tetraspanin detection was performed with αCD9, αCD63 or αCD81 antibody on milk EVs **(a)** and serum EVs **(b)**. Tetraspanin capture beads CD9, CD63 and CD81 are not included in this figure (Data are presented in Figure 5 and Supplementary Figure 5). Grey bars indicate non-significant detection compared to isotype control. a.u. = arbitrary unit.
